# Regulatory network analysis of *Dclk1* gene expression reveals a tuft cell-ILC2 axis that inhibits pancreatic tumor progression

**DOI:** 10.1101/2024.08.30.610508

**Authors:** Giovanni Valenti, Pasquale Laise, Ryota Takahashi, Feijing Wu, Tuo Ruan, Alessandro Vasciaveo, Zhengyu Jiang, Masaki Sunagawa, Moritz Middelhoff, Henrik Nienhüser, Na Fu, Ermanno Malagola, Yoku Hayakawa, Alina C. Iuga, Andrea Califano, Timothy C. Wang

## Abstract

*Dclk1* expression defines a rare population of cells in the normal pancreas whose frequency is increased at early stages of pancreatic tumorigenesis. The identity and the precise roles of *Dclk1* expressing cells in pancreas have been matter of debate, although evidence suggests their involvement in a number of key functions, including regeneration and neoplasia. We employed a recently developed Dclk1 reporter mouse model and single cell RNAseq analysis to define *Dclk1* expressing cells in normal pancreas and pancreatic neoplasia. In normal pancreas, *Dclk1* epithelial expression identifies subsets of ductal, islet and acinar cells. In pancreatic neoplasia, *Dclk1* expression identifies five epithelial cell populations, among which acinar-to-ductal metaplasia (ADM)-like cells and tuft-like cells are predominant. These two cell populations play opposing roles in pancreatic neoplasia, with Dclk1^+^ ADM-like cells sustaining tumor growth while Dclk1^+^ tuft-like cells restraining tumor progression. The differentiation of Kras mutant acinar cells into Dclk1^+^ tuft-like cells requires the activation of the transcription factor SPIB and is further supported by a cellular paracrine loop involving cancer group 2 innate lymphoid cells (ILC2) and cancer activated fibroblasts (CAFs) that provide IL13 and IL33, respectively. In turn, Dclk1^+^ tuft-like cells release angiotensinogen that plays protective roles against pancreatic neoplasia. Overall, our study provides novel insights on the biology of Dclk1^+^ cells in normal pancreas and unveils a protective axis against pancreatic neoplasia, involving CAFs, ILC2 and Dclk1^+^ tuft-like cells, which ultimately results in angiotensinogen release.

## Introduction

Pancreatic ductal adenocarcinoma (PDAC) remains one of the most lethal cancer types with a 5-year survival rate in the U.S. of 9%^1^. The genetic changes that contribute to the development of PDAC have been well described now, with KRAS mutated in 93% of patients^2^. Conversely, the debate on the cell of origin of PDAC is still ongoing. Pancreatic intraepithelial neoplasia (PanIN) represents the most common precursor lesion for pancreatic cancer^3^. PanINs comprise cells exhibiting features of ductal cells, which initially suggested the hypothesis of a pancreatic ductal cell of origin of PDAC^4^. However, accumulating evidence has pointed to pancreatic acinar cells as the most likely source of PDAC, given also their capacity to undergo acinar-to-ductal metaplasia (ADM) in response to injury, where they acquire features of ductal cells^5,6,7^.

While mathematical modelling indicates a uniform progenitor potential among all acinar cells, lineage tracing studies have revealed an intrinsic heterogeneity of acinar cells, with only minor subsets retaining the capacities to proliferative long term and to regenerate pancreas following injuries, therefore serving as putative progenitor cells^8,9,10,11,12^. To add further complexity to this scenario, a population of transit amplifying acinar progenitors that lack regenerative potential and are resistant to malignant transformation has also been recently reported^13^. Together, these findings have raised the question on whether all acinar cells can undergo ADM and therefore initiate PDAC, or whether such as capacity would be only reserved to specific subsets of acinar cells, such as those involved in pancreas regeneration following pancreatic injuries^14^.

The Doublecortin like kinase 1 (*Dclk1*) gene was initially reported to be expressed in normal pancreas by subsets of islet and ductal cells, but subsequently low levels of *Dclk1* expression were found also in a rare population of quiescent acinar cells that are dispensable under homeostatic conditions, but essential for pancreas regeneration^8,15^. Dclk1^+^ acinar cells exhibit a superior capacity to initiate pancreatic tumors in presence of *Kras* activating mutations and tissue injury^8^. An increased frequency of Dclk1^+^ cells has been described in early stages pancreatic neoplasia in diverse mouse models.

However the identity of these Dclk1^+^ cells has been matter of debate^16,17^. A number of studies have reported these Dclk1^+^ cells as putative cancer stem cells (CSCs) and potential cells of origin for PDAC^8,16,18^. Other studies have described these Dclk1^+^ cells as bone-fide tuft cells, a rare chemosensory cell type, which would play a tumor protective role in pancreatic neoplasia^17,19,20^.

Tuft cells are rare post-mitotic cells that are more typically found in the intestine and respiratory tract and can be identified by their unique morphology and expression of a specific array of markers, including DCLK1, TRPM5, POU2F3, CD24A^21,22,23,24^. Tuft cells were originally described as involved in sensing and integrating signals from the microenvironment, and in organizing appropriate responses in order to maintain tissue homeostasis^21,22^. In the intestine, tuft cells play central roles in initiating and coordinating immune responses to helminthic infections through an interaction loop with resident group 2 innate lymphoid cells (ILC2). Tuft cells sense metabolites produced by helminths and in response release IL25 that stimulates ILC2 to initiate the appropriate immune responses^25,26,27,28,29^. Activated ILC2 sustain this response by promoting tuft cell expansion via type 2 cytokines, such as IL4 and IL13, thereby creating a feedback loop that is crucial to resolve infections^25,26,27^. While this model is relevant to the gut in the context of helminthic infections, different cellular loops could be realized in other organs. For instance, IL33 has been reported as the primary cytokine responsible for recruiting and expanding ILC2 in other tissues, where it contributes to tissue regeneration and healing^30^.

ILC2 are present and often increased in tumor settings, but interestingly they have been associated with both pro-tumorigenic and anti-tumorigenic responses depending on the type and stage of cancer^31,32^. In PDAC, it has been recently shown that ILC2 are mostly anti-tumorigenic, as they infiltrate pancreatic tumors and activate dendritic and T CD8 cells^33^. While the focus of these studies has been primarily on the interactions of ILC2 with other immune cells, their potential effect on epithelial cells in the settings of neoplasia has not been fully explored.

To study *Dclk1* expressing cells more extensively, we employed a reporter mouse we recently generated (Dclk1-ZsGreen) and transcriptionally profiled Dclk1^+^ cells isolated from both normal pancreas and early pancreatic neoplasia by single cell RNA-seq. In normal pancreas, *Dclk1* expression identifies a heterogeneous cell population that includes subsets of ductal, islet and acinar cells. In early pancreatic neoplasia, *Dclk1* expression identifies two main cell populations, ADM-like and tuft-like cells, both arising from Kras mutant acinar cells through distinct differentiation programs. The two cell types play opposite roles in pancreatic neoplasia, as Dclk1^+^ ADM like cells sustain tumor growth while Dclk1^+^ tuft like cells restrain tumor progression. The protective function of Dclk1^+^ tuft-like cells is the results of a cellular paracrine loop that also involves ILC2 and CAFs and results in the release by these cells of angiotensinogen, which play important roles against pancreatic tumor progression. Overall, our study provides novel insights on Dclk1^+^ cells in normal pancreas and pancreatic neoplasia, and supports the emerging concept of the protective roles for tuft cells in neoplasia, highlighting also critical functions of angiotensinogen in this disease.

## Results

### *Dclk1* expression identifies subsets of ductal, endocrine and acinar cells in normal pancreas

To study *Dclk1* expressing cells in the normal pancreas, we used a Dclk1 reporter mouse model that we generated (Dclk1-DTR-ZsGreen), in which the gene of the fluorescent protein ZsGreen was placed downstream of the *Dclk1* gene regulatory elements in a BAC vector (Figure S1A). ZsGreen expressing cells in the pancreas of these mice stained positive for DCLK1 and expressed higher levels of *Dclk1* with respect to the other pancreatic epithelial cells, confirming that ZsGreen expression provided an accurate readout for *Dclk1* expression (Figures S1B-C). Epithelial ZsGreen^+^ cells in the normal pancreas accounted for approximately 0.3% of total pancreatic cells, and were distributed in a mosaic fashion in pancreas epithelium, with the majority localized to the ducts and minor fractions in islets and acini (Figures 1A-C). In support of this distribution pattern, ZsGreen^+^ cells were highly enriched within the ductal CD49f^+^CD133^+^ cell population as assessed by flow cytometry, and also exhibited higher expression of ductal markers over islet and acinar markers (Figures 1D-F, S1D)^34^.

**Figure 1.**
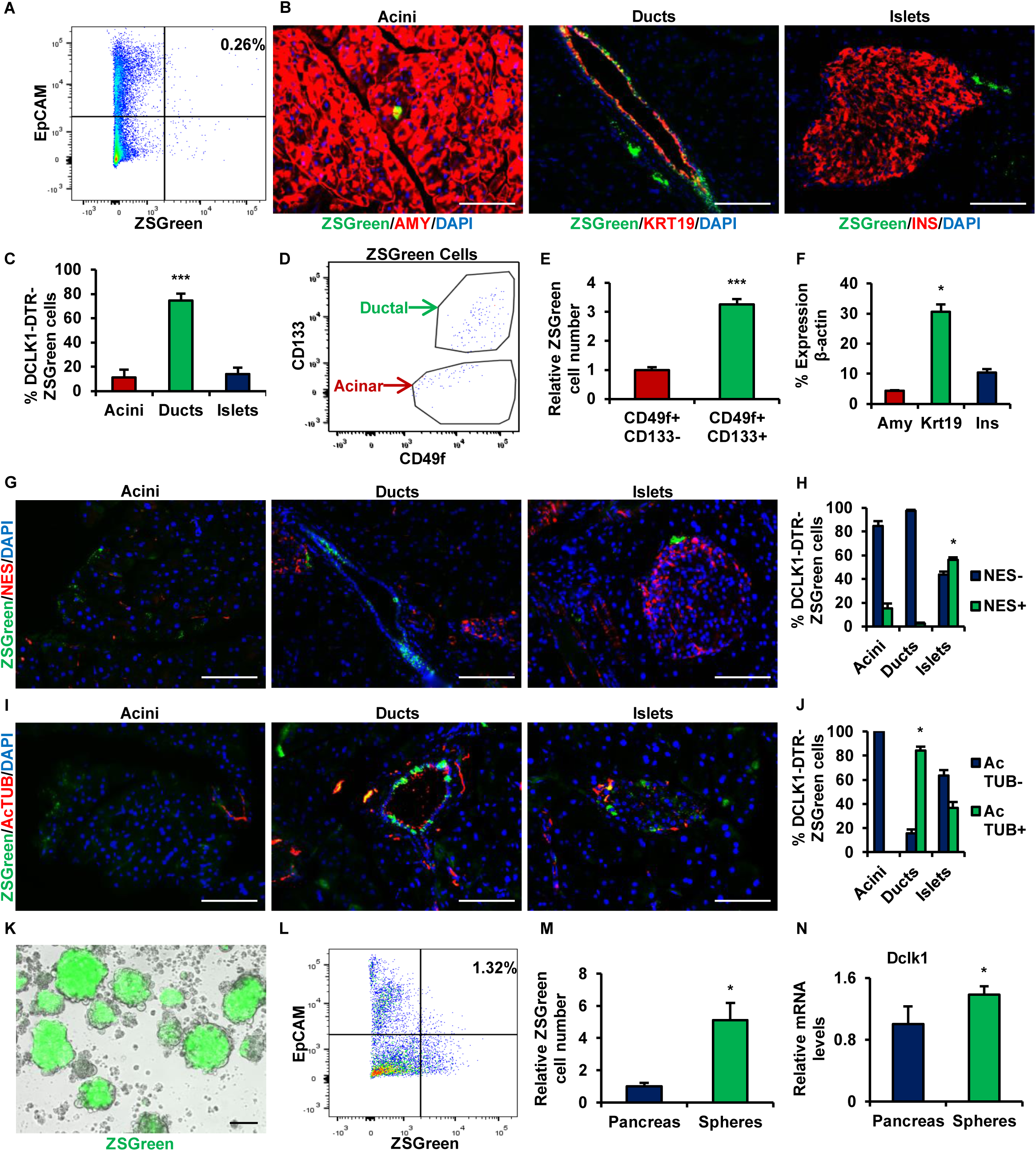
**(A)** Flow cytometry analysis of single cells isolated from normal pancreas of Dclk1-DTR-ZsGreen mice for EpCAM and ZsGreen. **(B)** Immunofluorescence of normal pancreas of Dclk1-DTR-ZsGreen mice for amylase (AMY), keratin 19 (KRT19) and Insulin (INS). **(C)** Quantification of ZsGreen cells localized in acini, ducts and islets based on immunofluorescence. **(D)** Flow cytometry analysis of ZsGreen^+^ cells isolated from normal pancreas of Dclk1-DTR-ZsGreen mice for CD133 and CD49f. **(E)** Quantification of relative ZsGreen cell number within CD49f^+^CD133^-^ and CD49f^+^CD133^-^ populations. **(F)** qRT-PCR for expression of *Amy*, *Krt19* and Ins in EpCAM^+^ZsGreen^+^ cells isolated from normal pancreas of Dclk1-DTR-ZsGreen mice. **(G-J)** Immunofluorescence of normal pancreas of Dclk1-DTR-ZsGreen mice for Nestin (NES) and Acetylated Tubulin (AcTUB) and related quantifications. **(K)** ZsGreen expression in non-adherent spheres grown from single cells isolated from normal pancreas of Dclk1-DTR-ZsGreen mice. **(L-M)** Flow cytometry analysis of non-adherent spheres for EpCAM and ZsGreen and related quantification. **(N)** qRT-PCR for *Dclk1* expression in non-adherent spheres. Scale bars: 100 μm. Means ± SD. ∗: p ≤ 0.05; ∗∗: p ≤ 0.01; ∗∗∗: p ≤ 0.001.

To understand whether distinct tissue localization within pancreas epithelium also correlated with specific cellular features, we evaluated the expression by ZsGreen cells of an array of differentiation markers (Figures 1G-J, S1E-H). The putative pancreatic progenitor cell marker Nestin was largely found in ZsGreen cells localized to the islets, with positive cells also detectable among cells localized to the acini; PDX1 was more highly expressed by cells localized to the ducts but also found in islet cells, although to a lesser extent. The tuft cell marker Acetylated Tubulin was largely found in ductal cells, although positive cells were also detected among cells localized to the islets; COX2 was expressed only by rare cells localized to the ducts.

ZsGreen cells and *Dclk1* expression were also enriched in spheres grown as non-adherent cultures from single cells isolated from normal pancreas, an assay known to enrich for progenitor cells, therefore supporting the possibility that a subset of Dclk1^+^ cells exhibited progenitor features (Figures 1K-N, S1I)^8,9,15^. Consistent with these findings, increased expression levels of other progenitor and ductal markers (*Nes1*, *Hes1*, *Krt19*) were also found in non-adherent spheres (Figures S1J-L).

To further characterize cell types identified by *Dclk1* expression in normal pancreas, we profiled 366 ZsGreen cells sorted from pancreas of Dclk1-DTR-ZsGreen mice by droplet-based scRNA-seq. Single-cell gene expression profiles were processed by the ARACNe-VIPER pipeline^35^. Briefly, the ARACNe algorithm was applied to generate a Dclk1-specific regulatory network and identify the regulatory target genes of 3770 regulatory proteins (RPs) including transcription factors (TFs), co-TFs and signaling molecules. Next, the VIPER algorithm was applied to the target genes of a given RP as inferred by ARACNe to infer its transcriptional activity. For simplicity, “protein activity” will be referred as the transcriptional activity of a given regulatory protein, as estimated by the VIPER algorithm.

Unsupervised clustering based on VIPER-inferred protein activity profiles partitioned cells into five clusters (Figures 2A-C). Analysis of differentially expressed genes revealed that three cell clusters corresponded to epithelial cells with features of pancreatic ductal cells (*Cd24a*, *Krt19*, *Prom1*, *Sox9*), islet cells (*Ins2*, *Trpm5*) and acinar cells (*Amy2a*, *Cela1*) (Figures 2D-F). The other two clusters were identified as immune cells (*Ptprc*) and stellate cells (*Vim*) (Figures S2A-B). ZsGreen cells exhibiting ductal features represented the majority of *Dclk1* expressing cells within pancreas epithelium.

**Figure 2.**
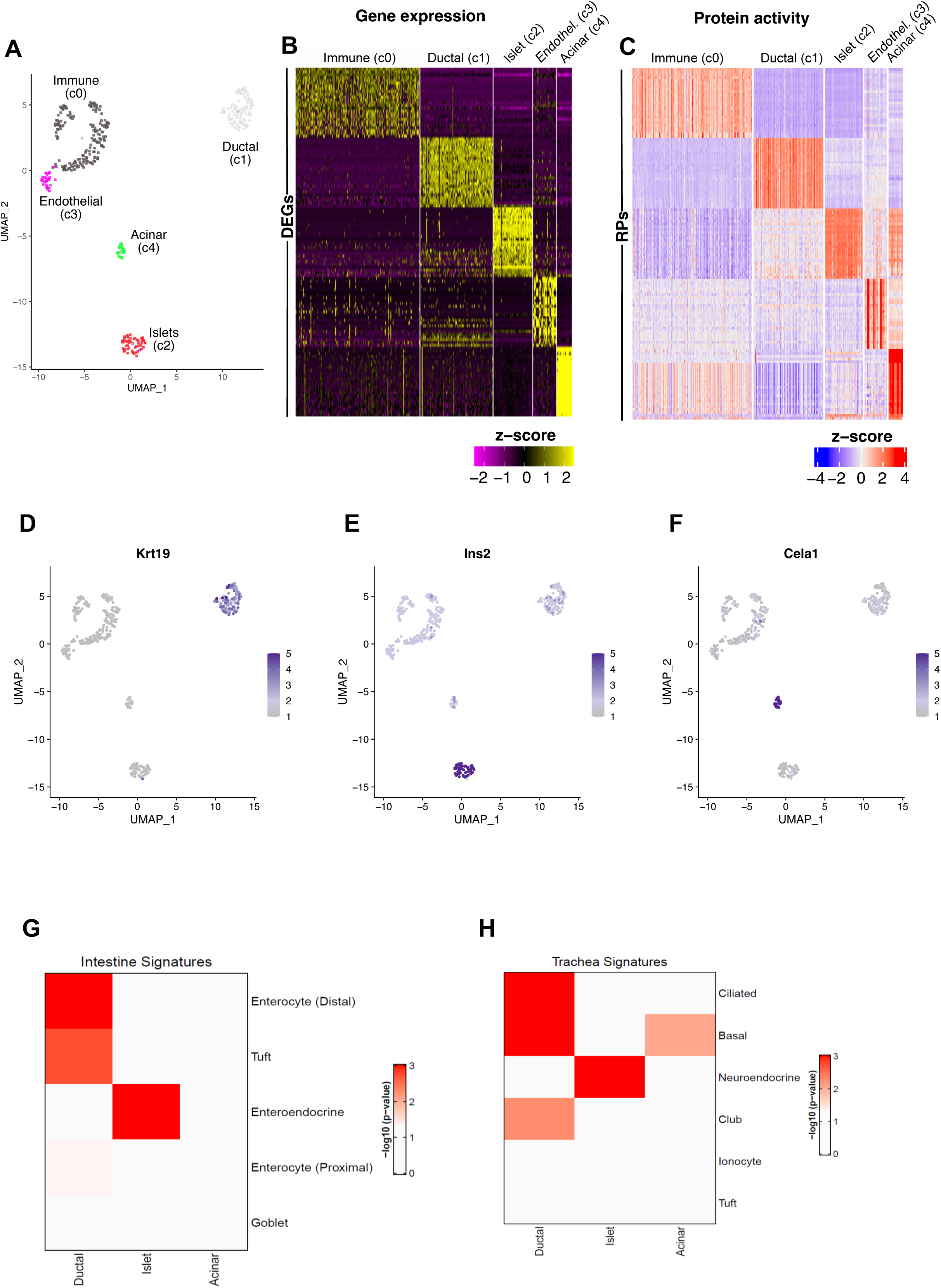
**(A)** UMAP based on VIPER activity showing different cell types identified by cluster analysis in Dclk1^+^ normal cells. **(B)** Heatmap showing differential expression between cell types in Dclk1^+^ normal cells. **(C)** Heatmap showing differential protein activity as inferred by VIPER algorithm, between cell types in Dclk1^+^ normal cells. **(D-F**) UMAP showing differential expression of marker genes across different cell types in Dclk1^+^ normal cells. **(G-H)** Heatmap showing the enrichment of publicly available gene expression signatures in Ductal, Islet and Acinar clusters identified in Dclk1^+^ normal cells. P-values were estimated by 1-tailed GSEA test with 1000 permutations.

To gain further insights into the potential functions of *Dclk1* expressing cells in pancreas epithelium, we studied their protein activity profiles. Dclk1^+^ ductal and islet protein did not show any major features that could be indicative of potential distinct functions from the bulk of pancreatic ductal and islet cells, respectively. In contrast, Dclk1^+^ acinar cell emerged to potentially have distinct functions with respect to the bulk of pancreatic acinar cells, given that we found high protein activity of several receptors for neurotrophic and axon guidance factors (*Adrb2*, *Ngfr*) as well as for immune signals (*Il4ra*, *Il17ra*), but the absence of other markers of pancreatic acinar cells (*Gata4*) (Figures S2E-H).

As a complementary strategy, we compared pancreatic Dclk1^+^ cell gene signatures with previous scRNA-seq signatures from intestine and trachea, given also that tuft cell clusters were well characterized in these other organs^23,24^ (Figures 2G-H). The Dclk1^+^ ductal cell signature was closely associated to those of enterocytes (distal) and tuft cells from the intestine as well as ciliated and basal cells from the trachea. The Dclk1^+^ islet cell signature was highly enriched within those of intestinal enteroendocrine cells and tracheal neuroendocrine cells. In contrast, the Dclk1^+^ pancreatic acinar cell signature did not associate with any cell types from both intestine and trachea. When performing this analysis against specific signatures for intestinal or tracheal tuft cells (tuft-1: neural tuft cells, tuft-2: immune tuft cells), only the Dclk1^+^ ductal cell signature displayed a significant enrichment (Figures S2I-J). Furthermore, several well-established tuft cell markers were included among the core genes of this enrichment analysis (*Cd24a*, *Dclk1*, *Krt18*, *Pou2f3*, *Ptgs1*).

Overall, our data show that *Dclk1* expression defines three cell populations in normal pancreas epithelium that are localized in ducts, islets and acini, and includes cells with potential tuft, neuroendocrine and progenitor cell features.

### Dclk1^+^ cells are expanded at early stage pancreatic neoplasia and display features of cancer stem cells or tuft cells

Next, we investigated *Dclk1* expressing cells in pancreatic neoplasia by crossing Dclk1-DTR-ZsGreen mice with two mouse models with different rates of disease progression. The first was a model of pancreatic intraepithelial neoplasia (PanIN) in which the expression of an activating Kras^G12D^ mutation could be targeted into acinar cells by the inducible recombinase Mist1-CreERT2 (Mist1-Kras)^17,36^. The second was a model of more advanced pancreatic tumors generated by introducing Kras^G12D^ and loss of function p53^R172H^ mutation in epithelial cells of the pancreas by the constitutive Pdx-Cre recombinase (KPC)^37,38^.

In both Mist1-Kras and KPC mice we observed a marked expansion of ZsGreen cells that was more pronounced in the Mist1-Kras model (Figures 3A-C, S3A). ZsGreen cell hyperplasia was largely localized to PanIN lesions and to a lesser extent in some normal ducts in both models, but it was attenuated in areas where the neoplasia was at the more advanced stages, which was especially evident in KPC mice. Despite the clear expansion of ZsGreen cells in pancreatic neoplasia, these cells were rarely positive for the proliferation marker KI-67, suggesting that this expansion was largely sustained by acinar cells that began to express *Dclk1* subsequently to KRAS activation, rather than proliferation of the resident pool of Dclk1^+^ acinar cells in normal pancreas (Figures 3D, S3B). In line with this model of expansion, lineage tracing studies carried out in Mist1-Kras-TdTomato-Dclk1-DTR-ZsGreen mice showed that a subset of acinar cells began upregulating both ZsGreen and DCLK1 as early as one week after KRAS activation, and more markedly over the course of the progression (Figures 3E, S3C).

**Figure 3.**
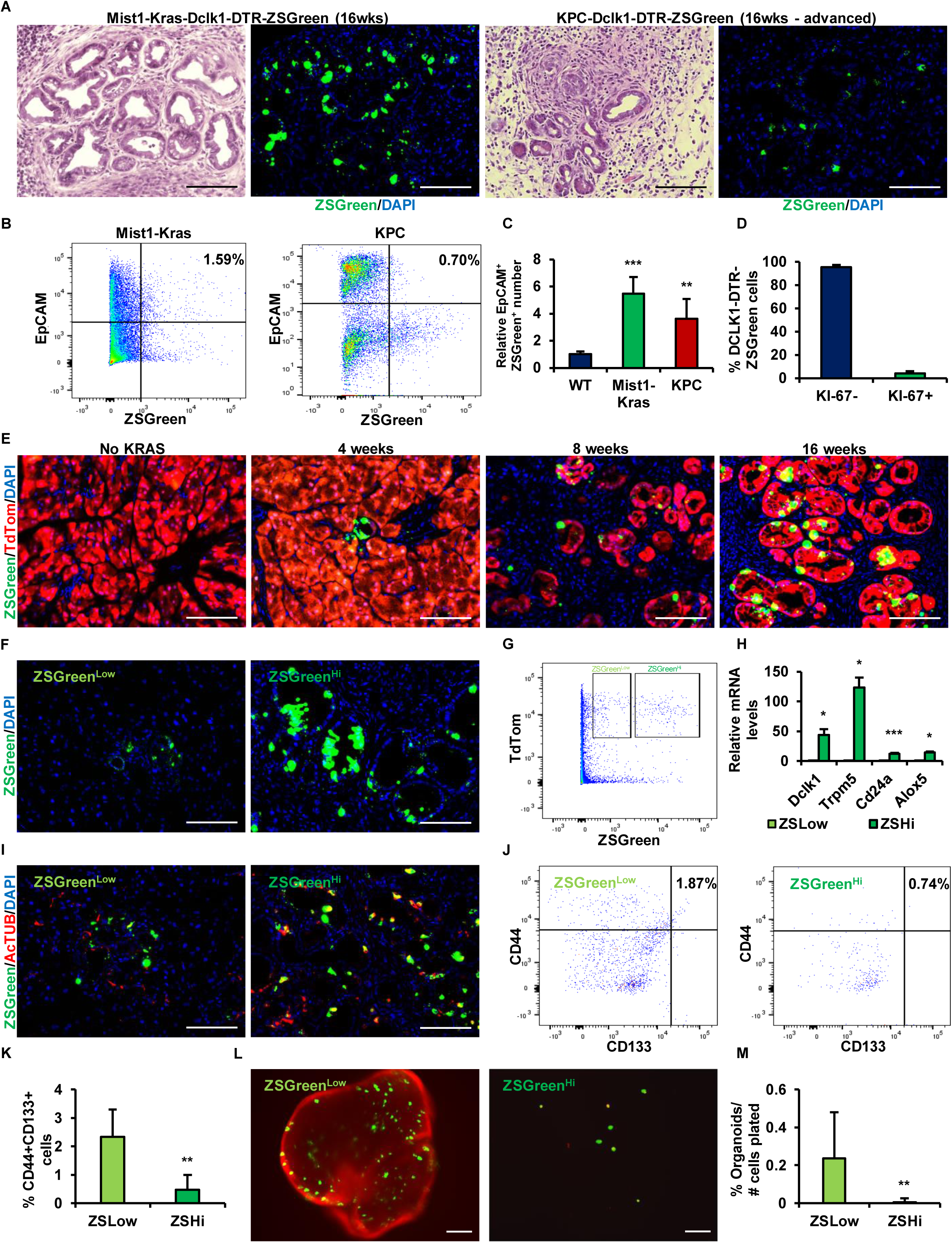
**(A)** Hematoxylin and Eosin staining (H&E) of pancreas from Mist1-Kras-Dclk1-DTR-ZsGreen mice and KPC-Dclk1-DTR-ZsGreen mice (area with advanced neoplasia) at 16 wks. **(B-C)** Flow cytometry analysis of single cells isolated from Mist1-Kras-Dclk1-DTR-ZsGreen mice and KPC-Dclk1-DTR-ZsGreen mice at 16 wks for EpCAM and ZsGreen and related quantification. **(D)** Quantification of ZsGreen^+^ cells expressing KI-67 based on immunofluorescence. **(E)** Immunofluorescence of pancreas of Mist1-Kras-Dclk1-ZsGreen-Rosa26-TdTomato mice not tamoxifen induced (No KRAS) and 4 wks, 8 wks and 16 wks after tamoxifen administration for ZsGreen and TdTomato (TdTom). **(F)** Immunofluorescence of pancreas of Mist1-Kras-Dclk1-ZsGreen showing ZsGreen cells with low (ZsGreen^Low^) or high expression (ZsGreen^Hi^). **(G)** Flow cytometry analysis of single cells isolated from pancreas of Mist1-Kras-Dclk1-DTR-ZsGreen-Rosa26-TdTomato mice for TdTom and ZsGreen showing the two populations of TdTomato^+^ZsGreen^low^ and TdTomato^+^ZsGreen^hi^ cells. **(H)** qRT-PCR for the expression of *Dclk1*, *Trpm5*, *Cd24a* and *Alox5* in isolated TdTomato^+^ZsGreen^low^ and TdTomato^+^ZsGreen^hi^ cells. **(I)** Immunofluorescence of pancreas of Mist1-Kras-Dclk1-ZsGreen mice for AcTUB in ZsGreen^low^ and ZsGreen^hi^ cells. **(J-K)** Flow cytometry analysis of ZsGreen^low^ and ZsGreen^hi^ cells for CD133 and CD44 and related quantification. **(L-M)** Organoids generated from isolated TdTomato^+^ZsGreen^low^ and TdTomato^+^ZsGreen^hi^ cells and related quantification. Scale bars: 100 μm. Means ± SD. ∗: p ≤ 0.05; ∗∗: p ≤ 0.01; ∗∗∗: p ≤ 0.001.

We observed that acinar cells expressed ZsGreen at variable levels within PanIN lesions, with some showing low levels (ZsGreen^Low^) while others high levels (ZsGreen^Hi^) of expression (Figure 3F). Distinct ZsGreen^Low^ and ZsGreen^Hi^ cell populations could be also discriminated by flow cytometry (Figure 3G). ZsGreen expression levels correlated well with the levels of *Dclk1* expression, as confirmed by qPCR analysis (Figure 3H). Given that *Dclk1* expression was reported to identify either cancer stem cells or tuft cells in pancreatic neoplasia by different studies, we next looked whether the different levels of *Dclk1* expression could discriminate between these two conditions. ZsGreen^Low^ cells exhibited no tuft cell morphology, low expression of tuft cell markers and were enriched for CD44^+^CD133^+^ surface markers, which are characteristic of cancer stem cells (Figures 3H-J, S3D-E)^39^. In contrast, ZsGreen^Hi^ cells showed tuft cell morphology, higher expression of tuft cell markers and lower fraction of CD44^+^CD133^+^ (Figures 3H-J, S3D-E). We therefore tested whether these differences in terms of markers also reflected differences at the functional levels by growing organoid cultures from single ZsGreen^Low^ and ZsGreen^Hi^ cells sorted from Mist1-Kras mice. Remarkably, ZsGreen^Low^ cells robustly generated organoids *in vitro*, whereas ZsGreen^Hi^ cells lacked of this capacity and showed no expansion as they remained as single cells in cultures (Figures 3L-M).

Overall, these results show that *Dclk1* expression by acinar cells is an early event after Kras activation, which results in the generation of at least two distinct cell populations, including one with stemness features (*Dclk1*/ZsGreen^Low^) and another resembling tuft cells (*Dclk1*/ZsGreen^Hi^).

### *Dclk1* expression defines at the molecular levels two main cell types in pancreatic neoplasia representing ADM-like cells and tuft-like cells

In order to further dissect the identity of Dclk1^+^ cells in early pancreatic neoplasia and to understand the mechanisms underlying their generation, we performed another scRNA-seq analysis. Specifically, we used a droplet-based scRNA-seq to profile 6301 Dclk1^+^ cells isolated from Mist1-Kras-TdTomato-Dclk1-DTR-ZsGreen mice at 2 weeks and 16 weeks after KRAS activation. TdTomato was used to distinguish Dclk1^+^ cells generated by *Kras* mutant acinar cells from other Dclk1^+^ cells, given that TdTomato provided readout for KRAS activation. Normal Dclk1^+^ cells isolated from pancreas of Dclk1-DTR-ZsGreen mice were also included in the analysis as additional internal control. Gene expression data were filtered for low quality cells and processed using the VIPER algorithm^35^. Cluster analysis based on inferred protein activity profiles showed higher robustness with respect to the same cluster analysis performed on gene expression data as assessed by silhouette analysis, which provided the rationale for relying on protein activity-based clusters (Figures S4A).

VIPER-based cluster analysis identified nine distinct clusters using the Louvain algorithm (as implemented in the Seurat package) and with the resolution parameter optimized by silhouette analysis (Figures 4A-C). TdTomato^+^ Dclk1^+^ cells from Mist1-Kras mice at 2 wks after KRAS activation showed partial overlap with Dclk1^+^ cells from normal mice and TdTomato^-^ Dclk1^+^ from Mist1-Kras mice, while TdTomato^+^ Dclk1^+^ cells from Mist1-Kras mice at 16 wks after KRAS activation partitioned into entirely distinct clusters (Figure S4B). *Dclk1* expression levels were variable among all the clusters, but markedly higher in TdTomato^+^ Dclk1^+^ cells at 16 wks after KRAS activation (Figures 4D-E). Moreover, *Dclk1* expression levels were negatively correlated with the levels of KRAS protein activity in profiled cells, as also confirmed by linear regression (Figures S4C-D).

**Figure 4.**
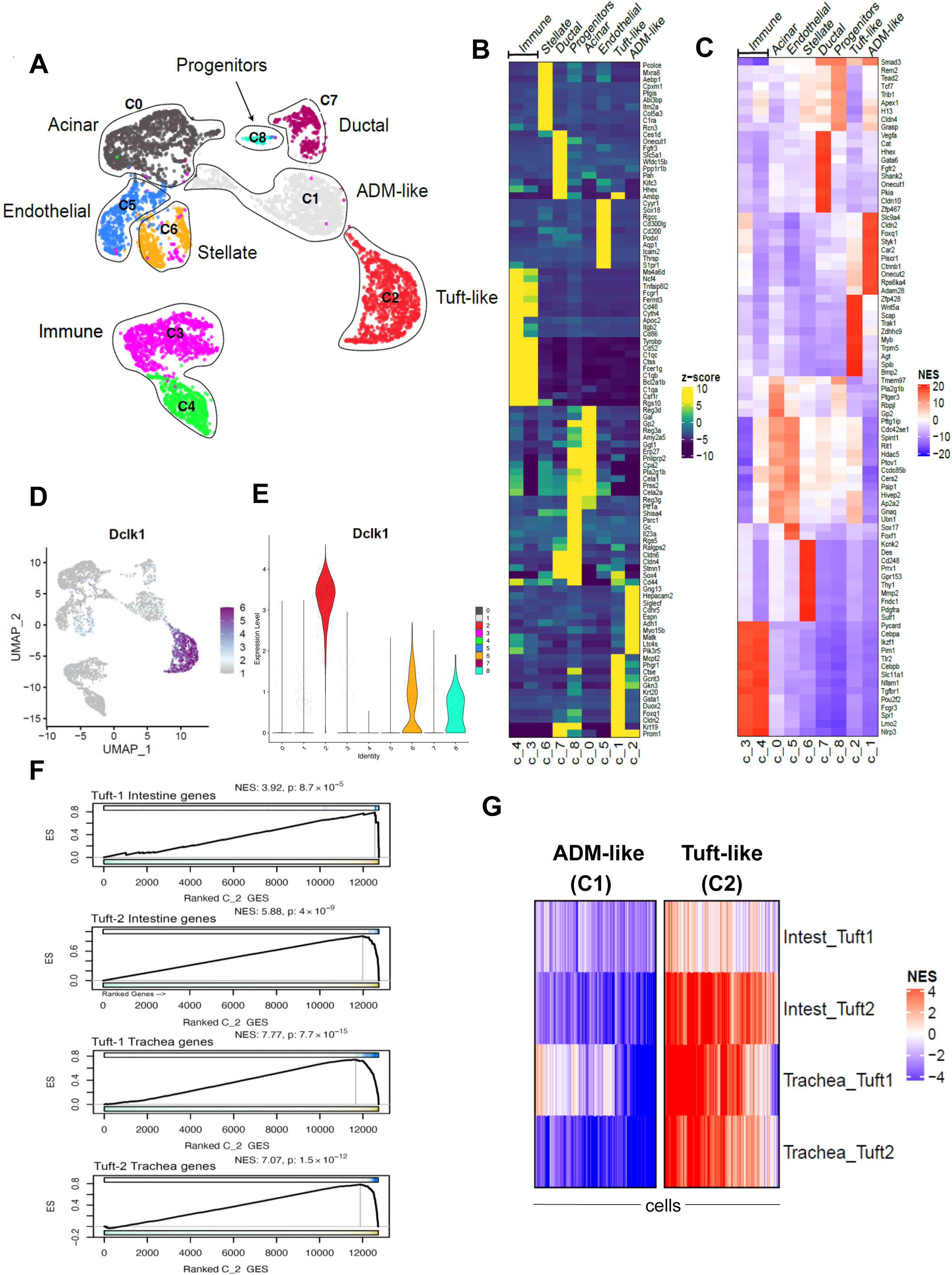
**(A)** UMAP based on VIPER activity showing different cell types identified by cluster analysis, including Dclk1^+^ normal cells and Dclk1^+^ cells after Kras activation at 2 weeks and 16 weeks. **(B)** Heatmap showing differential expression across cell types. **(C)** Heatmap showing differential protein activity across cell types. **(D)** UMAP showing *Dclk1* expression. **(E)** Violin plots showing *Dclk1* expression across clusters/cell types. **(F)** GSEA plots showing the enrichment of Tuft gene expression signatures derived from intestine and trachea in the gene expression signature of tuft-like cells cluster (C2). Normalized enrichment score (NES) and p-values were estimated by 1-tailed GSEA test with 1000 permutations. **(G)** Heatmaps showing the enrichment of Tuft gene expression signatures derived from intestine and trachea in ADM-like (C1) and Tuft-like (C2) cells. Normalized enrichment scores were estimated by 1-tailed GSEA test with 1000 permutations.

Gene expression analysis using cell type-specific markers, identified five clusters of epithelial cells and four clusters of stromal cells, including two of immune cells, one of endothelial cells and one of stellate cells (Figures S4E-I). Among the epithelial clusters, one cluster (C7) was enriched for normal Dclk1^+^ cells and TdTomato^-^Dclk1^+^ cells at both 2 wks and 16 wks; two clusters (C0 and C8) were enriched for TdTomato^+^Dclk1^+^ cells at 2 wks, TdTomato^-^Dclk1^+^ cells at 2 wks and normal Dclk1^+^ cells; two clusters (C1 and C2) were enriched for TdTomato^+^Dclk1^+^ cells at 16 wks. C7 (Ductal) included cells showing low *Dclk1* expression and high KRAS activity that largely exhibited features of Dclk1^+^ ductal cells found in normal pancreas (*Hes1*, *Krt19*, *Sox4*, *Spp1*), sharing with them approximately 80% of differentially activated proteins and approximately 75% of markers, with differences accounting for genes involved in chromatin remodeling, inflammatory responses and hypoxia. C0 (Acinar) was a large cluster of cells showing low *Dclk1* expression and low KRAS activity that overall exhibited features of normal Dclk1^+^ acinar cells (*Amy2a5*, *Cela1*, *Cela2a*, *Prss2*), sharing with them approximately 95% of differentially activated proteins and approximately 85% of markers. C8 (Progenitors) was a small cluster of cells characterized by high *Dclk1* expression and high KRAS activity that showed features of putative progenitor cells, as highlighted by the expression of acinar, ductal and progenitor markers (*Prss2*, *Ptf1a*, *Sox4*, *Stmn1*). C1 (ADM-like) was a large cluster of cells showing low *Dclk1* expression and high KRAS activity and characterized by the expression of ADM markers (*Krt19*, *Sox4*) as well as CSC markers (*Cd44*, *Prom1*) (Figures S4J-K). C2 (Tuft-like) was a large cluster of cells characterized by high *Dclk1* expression and low KRAS activity that displayed features of tuft cells (*Cd24a*, *Pou2f3*, *Siglecf*, *Trpm5*), as also confirmed by the enrichment of its signature within the signatures of both immune and neural tuft cells from the intestine and the trachea (Figures 4F-G, S4L-M).

We next analyzed the four clusters defined by TdTomato^+^Dclk1^+^ cells at both 2 wks and 16 wks (acinar, progenitor, ADM-like, tuft-like) in order to infer potential differentiation trajectories followed by *Kras* mutant Dclk1^+^ acinar cells during the progression of pancreatic neoplasia (Figures 5A-B). Pseudo-trajectory analysis as inferred in the PCA and principal curve analysis revealed two potential differentiation trajectories that independently originated from Kras mutant Dclk1^+^ acinar cells and resulted in the generation of either ADM-like cells or tuft-like cells. The Dclk1^+^ progenitor cell cluster appeared not involved in any of these differentiation trajectories (Figure S5A-B).

**Figure 5.**
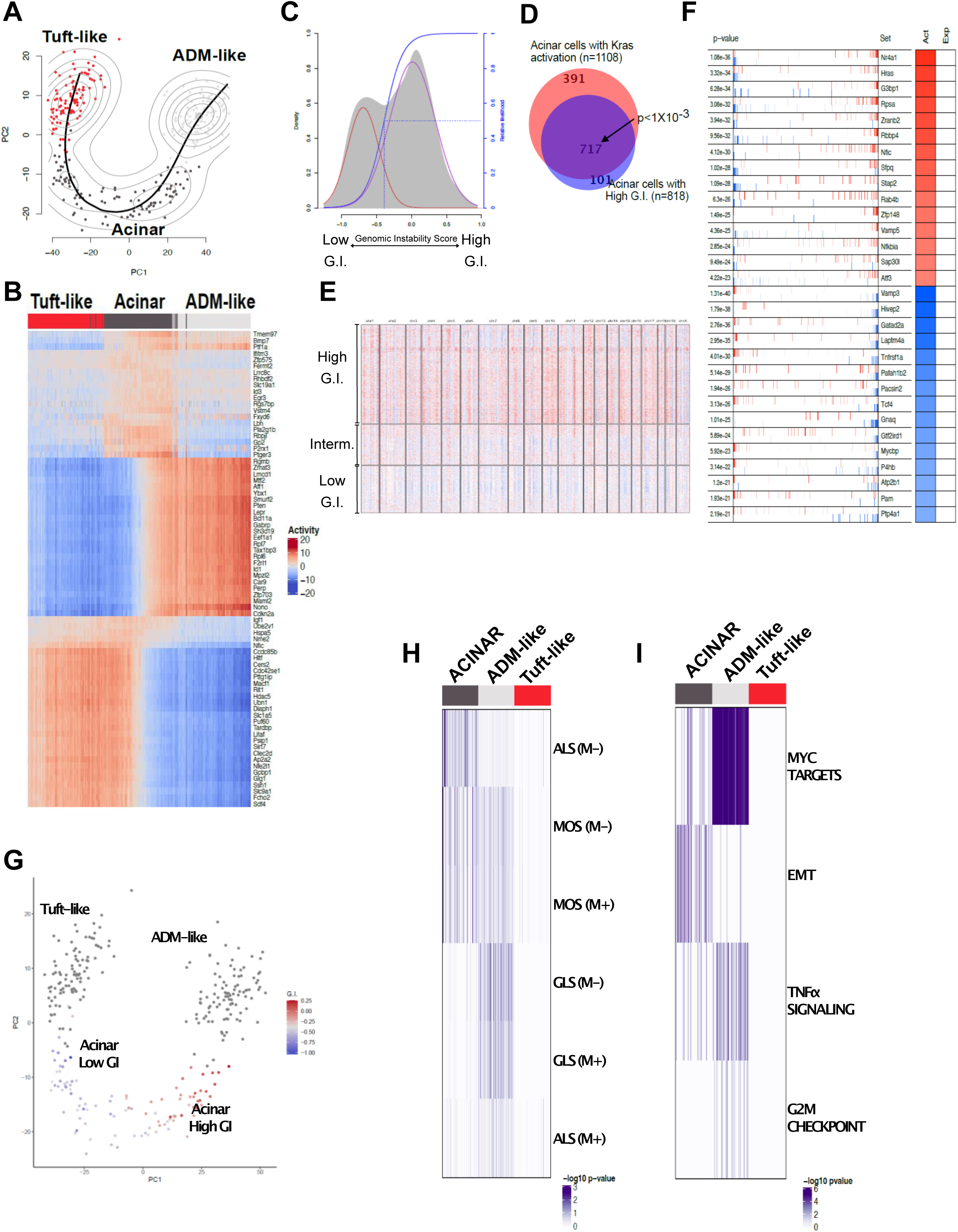
**(A)** PCA plot showing pseudo trajectory analysis across Tuft-like, Acinar and ADM-like cells. Pseudo trajectory was computed by principal curve analysis on the most representative cells (n=100) of each cluster/cell type selected by silhouette analysis. **(B)** Heatmap showing top differential activated proteins of Tuft-like, Acinar and ADM-like cells. Columns represent cells and are sorted based on the pseudo trajectory. Activity score is expressed as NES. **(C)** Density plot showing distribution of genomic instability score (GIS) inferred from gene expression profiles of Acinar cells with Kras activation at 2 weeks, compared to normal acinar cells. **(D)** Venn diagram plot showing the overlap between Acinar cells with Kras activation after 2 weeks and Acinar cells with high genomic instability as inferred by CNV analysis of single cell gene expression profiles. This plot shows that 65% (717/1108) of acinar cells with Kras activation after 2 weeks show higher genomic instability compared to normal cells. P-value was estimated by Fisher test as implemented in the ‘GenOverlap’ package. **(E)** Heatmap showing CNV inferred from single cell gene expression profiles of acinar cells with Kras activation after 2 weeks compared to normal acinar cells. Columns represent genomic regions and rows represent cells. Red indicates amplification, blue indicates deletion. **(F)** VIPER plot showing the top differentially activated proteins between acinar cells with Kras activation after 2 weeks and high GIS compared to normal acinar cells. **(G)** PCA plot showing the GIS of the acinar cells after 2 weeks of Kras activation including Tuft-like, Acinar and ADM-like cells. GIS correlates with pseudo trajectory analysis (Fig.5A) with the subset of acinar cells primed towards tuft-like cells showing lower GIS and acinar cells primed towards ADM-like cells showing higher GIS. **(H)** Heatmap showing the enrichment of the PDAC human subtypes in Tuft-like, Acinar and ADM-like cells. The enrichment was estimated by 1-tailed aREA test (as implemented in the VIPER package) with the top differentially activated proteins (n=50) of each human subtype in protein activity signature of Tuft-like, Acinar and ADM-like cells. Mouse gene names were converted to human gene names using the homologene package available on CRAN (https://cran.r-project.org/)^45^. **(I)** Heatmap showing the enrichment of the most enriched hallmarks gene sets in Tuft-like, Acinar and ADM-like cells. The enrichment was estimated by 1-tailed aREA test.

We hypothesized that while the short term 2 wks KRAS activation period was not sufficient to fully reprogram acinar cells into a different phenotypic state (ADM-like, tuft-like), this time window could be long enough to induce early phenotypic changes in Dclk1^+^ acinar cells towards neoplastic transformation, and more specifically could affect copy number variation (CNV)^39,40,41,42,43,44^. We therefore performed CNV analysis based on single cell gene expression profiles and inferred the genomic instability score of each individual cell among the 1108 Dclk1^+^ cells included in the Dclk1^+^ acinar cell cluster^45^. Using normal Dclk1^+^ acinar cells as a reference, we identified a population of 818 acinar cells (74%) showing increased CNV and high genomic instability score, which included a subset of 717 cells (88%) showing KRAS activation (Figures 5C-E). Differential protein activity analysis between Dclk1^+^ acinar cells with high and low genomic instability scores highlighted marked differences in proteins involved in TGF-α/NF-κB signaling, p53 pathway, MYC targets, G2M checkpoint and hypoxia (Figure 5F). Pseudo-trajectory analysis, as inferred in the PCA and principal curve analysis, revealed that while Dclk1^+^ acinar cells with high genomic instability scores generated ADM-like cells, Dclk1^+^ acinar cells with low genomic instability scores differentiated into tuft-like cells (Figure 5G). Collectively, these results indicate that Dclk1^+^ acinar cells are primed towards new phenotypic states as early as 2 wks after KRAS activation, and they subsequently reprogram into either ADM-like cells or tuft-like cells depending on their genomic instability (Figure S5C).

We compared our acinar, ADM-like and tuft cell signatures to signatures of human PDAC cell lineages identified in a meta-analysis conducted on scRNA-seq profiles from three public datasets (Figure 5H)^45^. The Dclk1^+^ acinar cell signature was significantly enriched in the ADM (ALS) and morphogenic (MOS) states while the Dclk1^+^ ADM-like signature was enriched in the gastrointestinal-lineage (GLS) and MOS states. The Dclk1^+^ tuft-like cell signature was not enriched in any PDAC cell lineages, supporting the notion of the absence of Dclk1^+^ tuft cells in advanced pancreatic tumors. Gene set enrichment analysis further revealed that the acinar cell signature was enriched for proteins involved in epithelial-to-mesenchymal transition (EMT), while the ADM-like cell signature was enriched for MYC targets and proteins involved in TGF-α/NF-κB signaling and G2M checkpoint (Figures 5I).

### SPIB as key transcription factor involved in tuft cell differentiation in pancreatic neoplasia

We were specifically interested in understanding better the potential genetic programs controlling the generation of Dclk1^+^ tuft-like cells at early pancreatic neoplasia. To this aim, we looked in more detail at transcription factors (TFs) that were selectively activated along the tuft cell differentiation trajectory. Of the 95 transcription factors included in the Dclk1^+^ tuft-like cell signature, several members of the ETS family displayed high protein activity (EHF, ELF4, ETV1, ETV6, SPDEF, SPIB) (Table S1). Among these, SPIB appeared to be of particular interest as its protein activity could be detected in Dclk1^+^ acinar cells as early as 2 wks after KRAS activation, it was the transcription factor showing the highest protein activity in Dclk1^+^ tuft-like cells, and it was also included in tuft cell gene signatures of other organs (Figure S5D)^21,23,24,28^. Consistent with a potential role in pancreatic neoplasia, *Spib* expression was increased in pancreas epithelium of Mist1-Kras mice with respect to normal pancreas epithelium, and selectively in Dclk1^+^ cells (Figures S5E-G). The gene network for SPIB *in vivo*, as inferred from our scRNA-seq data (see methods), included *Dclk1* as well as other well-established tuft cell markers, like *Cd24a* and *Sox9*, supporting the possibility that SPIB could regulate their expression (Figure S5H). To better identify the genetic programs controlled by SPIB in pancreatic neoplasia, we performed a RNA-seq analysis of pancreatic organoids grown from single cells obtained from normal murine pancreas, in which we overexpressed SPIB as well as mutant KRAS. SPIB overexpressing organoids displayed increased expression of *Dclk1* as well as other tuft cell markers, including *Cd24a* and *Sox9*, confirming its role in Dclk1^+^ tuft-like cell differentiation (Figures S5I-K). Gene set enrichment analysis revealed that SPIB mainly controlled the expression of genes involved in immunity and cytokine signaling.

### Dclk1^+^ cell hyperplasia is sustained by type 2 immunity in pancreatic tumors

KRAS activation in pancreatic organoids resulted in reduced ZsGreen cell number and *Dclk1* expression, suggesting that Dclk1^+^ cell hyperplasia at early stages pancreatic tumors was dependent on a combination of KRAS activation in acinar cells and exogenous signals from the microenvironment (Figures S6A-B).

Tuft cell hyperplasia was reported to be sustained by type 2 cytokines in the intestine during helminthic infections^25,26,27^. We therefore investigated whether IL4/IL13 signaling could play roles in promoting Dclk1^+^ tuft-like cell hyperplasia in pancreatic tumors. Our scRNA-seq data revealed IL4RA protein activity in Dclk1^+^ acinar cells at 2 wks after KRAS activation as well as in ADM-like and tuft-like cells, but not in progenitors or ductal cells (Figure 6A). In addition, *Il4ra* was more highly expressed by Dclk1^+^ cells with respect to other pancreatic epithelial cells of Mist1-Kras mice, consistent with its potential role in Dclk1^+^ cell differentiation (Figures S6C-D). In terms of IL4RA ligands, we found a marked increase in the frequency of IL13-producing cells in Mist1-Kras mice but not of IL4 (Figures 6B-C, S6E-F). This correlated with overall higher levels of *Il13* expression over *Il4* (Figure S6G).

**Figure 6.**
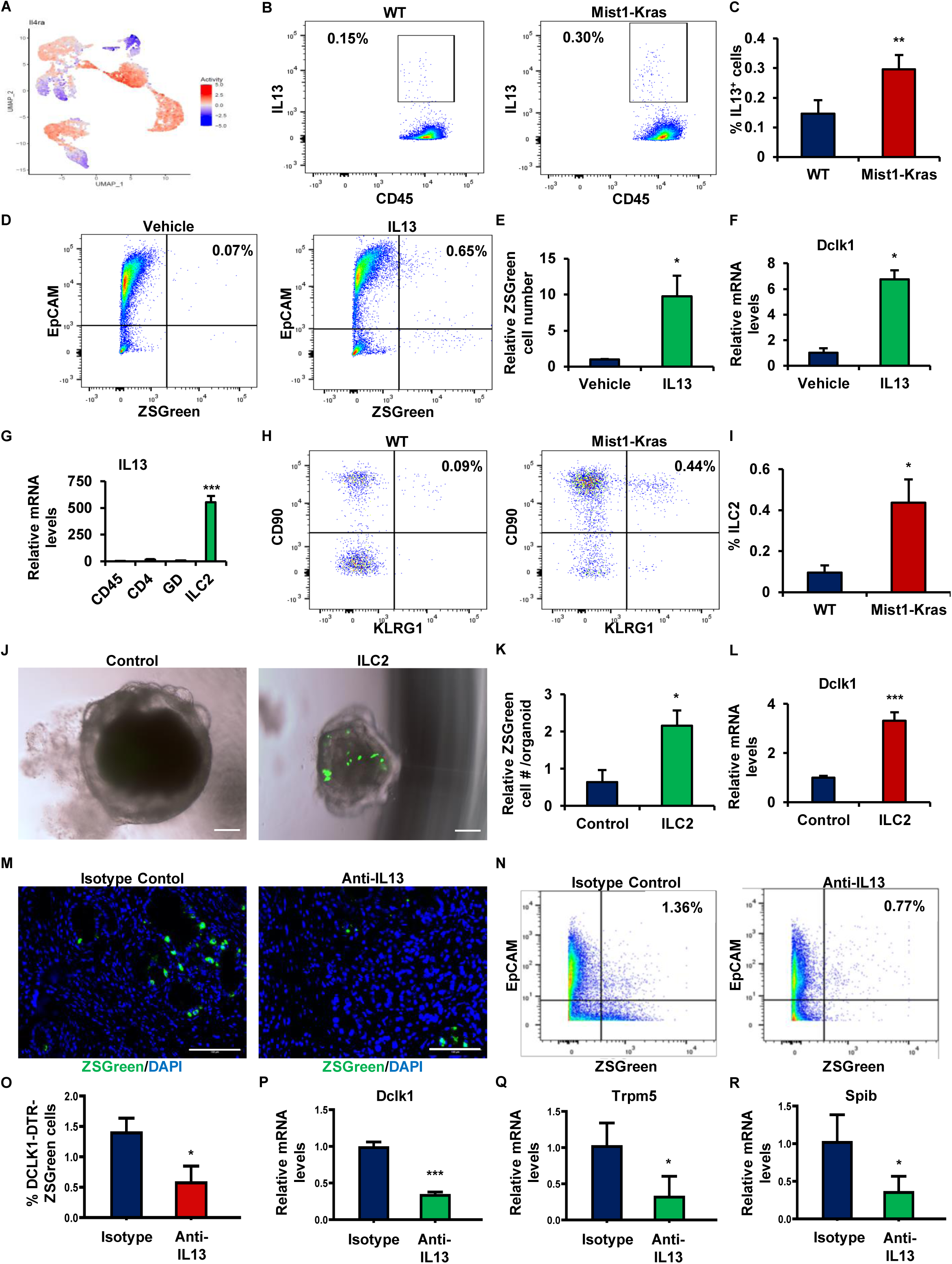
**(A)** UMAP showing VIPER-inferred differential protein activity of IL4RA in Dclk1^+^ cells. **(B-C)** Flow cytometry analysis for IL13 expression in immune cells (CD45) isolated from normal pancreas and pancreas from Mist1-Kras mice and related quantification. **(D-E)** Flow cytometry analysis for EpCAM and ZsGreen of organoids grown from single cells isolated from pancreas of Kras-Dclk1-DTR-ZsGreen in the presence of vehicle or IL13 and related quantification. **(F)** qRT-PCR for the expression of *Dclk1* in organoids grown in the presence of vehicle or IL13. **(G)** qRT-PCR for the expression of *Il13* in different immune cell populations in pancreas from Mist1-Kras mice: CD45^+^ immune cells, CD4^+^ T cells, γδ T cells and ILC2 (Lin^-^CD45^+^CD127^+^CD90^+^KLRG1^+^). **(H-I)** Flow cytometry analysis for ILC2 in normal pancreas and pancreas from Mist1-Kras mice and related quantification. **(J-K)** ZsGreen expression in organoids grown from single cells isolated from pancreas of Kras-Dclk1-DTR-ZsGreen mice in control medium or in the presence of ILC2 and related quantification. **(L)** qRT-PCR for the expression of *Dclk1* in organoids grown in the presence of control medium or ILC2. **(M)** Immunofluorescence for ZsGreen of pancreas from Mist1-Kras-Dclk1-DTR-ZsGreen mice treated with control IGG or anti-IL13 blocking antibody for 8 wks. **(N-O)** Flow cytometry analysis for EpCAM and ZsGreen of single cells isolated from pancreas of Mist1-Kras-Dclk1-DTR-ZsGreen mice treated with control IGG or anti-IL13 blocking and related quantification. **(P-R)** qRT-PCR for the expression of *Dclk1*, *Cd24a* and *Spib* in pancreas of Mist1-Kras-Dclk1-DTR-ZsGreen mice treated with control IGG or anti-IL13 blocking. Scale bars: 100 μm. Means ± SD. ∗: p ≤ 0.05; ∗∗: p ≤ 0.01; ∗∗∗: p ≤ 0.001.

We therefore tested whether IL13 could sustain Dclk1^+^ cell hyperplasia *in vitro* in pancreatic organoids grown from single cells isolated from Kras-Dclk1-DTR-ZsGreen mice. IL13 administration resulted in a significant increase in both Dclk1^+^ cell number and *Dclk1* expression, although overall organoid number was not affected (Figures 6D-F, S6H-I). We screened several immune cell populations isolated from Mist1-Kras mice for *Il13* expression, and identified ILC2 as the main cellular source of this cytokine (Figure 6G). Of note, ILC2 expressed significantly higher levels of *Il13* than *Il4* (Figure S6J). ILC2 frequency was increased in Mist1-Kras mice with respect to normal pancreas (Figures 6H-I). ILC2 isolated from Mist1-Kras mice sustained both Dclk1^+^ cell expansion and *Dclk1* expression *in vitro* in pancreatic organoids, therefore recapitulating the effects of IL13 administration (Figures 6J-L).

We looked at potential signals responsible for ILC2 activation in Mist1-Kras mice and focused primarily on alarmins (*Il25*, *Il33*, *Tslp*), which are known as potent ILC2 activators^46,47^. We found markedly higher levels of IL33 over the other alarmins in Mist1-Kras mice, suggesting that this cytokine could contribute to ILC2 activation in this model (Figures S6K-M). In line with this possibility, stimulation of ILC2 with IL33 resulted in a marked increase in *Il13* expression (Figure S6N). We investigated the cellular source of IL33 by screening its expression in several cell populations isolated from Mist1-Kras mice and identified cancer associated fibroblasts (CAFs) as a main cellular source of this cytokine (Figures S6O-P).

We next tested whether blocking IL13 signaling could diminish the generation of Dclk1^+^ cells in early pancreatic neoplasia in Mist1-Kras mice. Chronic administration over an eight week period of blocking antibody against IL13 resulted in both reduced Dclk1^+^ cells and *Dclk1* expression (Figures 6M-P). Furthermore, mice treated with the anti-IL13 antibody also displayed reduced expression levels of several tuft cell markers including *Cd24a*, *Sox9*, *Spib*, *Trpm5* as well as Acetylated Tubulin, consistent with an impaired differentiation of Dclk1^+^ acinar cells into tuft-like cells (Figures 6Q-R, S6Q-S).

Overall, these data show that IL13 produced by ILC2 likely plays a major role in Dclk1^+^ cell hyperplasia in early pancreatic neoplasia and blocking this signaling also results in reduced Dclk1^+^ tuft-like cells.

### Angiotensinogen is a key signal provided by Dclk1^+^ tuft cells to restrain the progression of pancreatic tumors

To uncover the functional roles of Dclk1^+^ tuft-like cells in pancreatic tumorigenesis, we screened our scRNA-seq data for secreted proteins expressed by these cells. Dclk1^+^ tuft-like cells expressed a limited number of ligands (Table S2). Among these, we found Angiotensinogen (*Agt*) of particular interest, given that it was the one showing the highest expression in Dclk1^+^ tuft-like cells (Figures 7A). In addition, we found that the *Agt* gene was also included in other intestinal tuft cell gene signatures, which further supported the possibility of its relevant roles for tuft cell function^21,23^. Angiotensinogen is the precursor of numerous signaling peptides originating from its proteolytic cleavage, among which angiotensin II is best known for its roles in the renin-angiotensin-aldosterone-system (RAAS) to regulate systemic blood pressure, although potential roles in cancer have been reported^50,51,52,53,54^.

**Figure 7.**
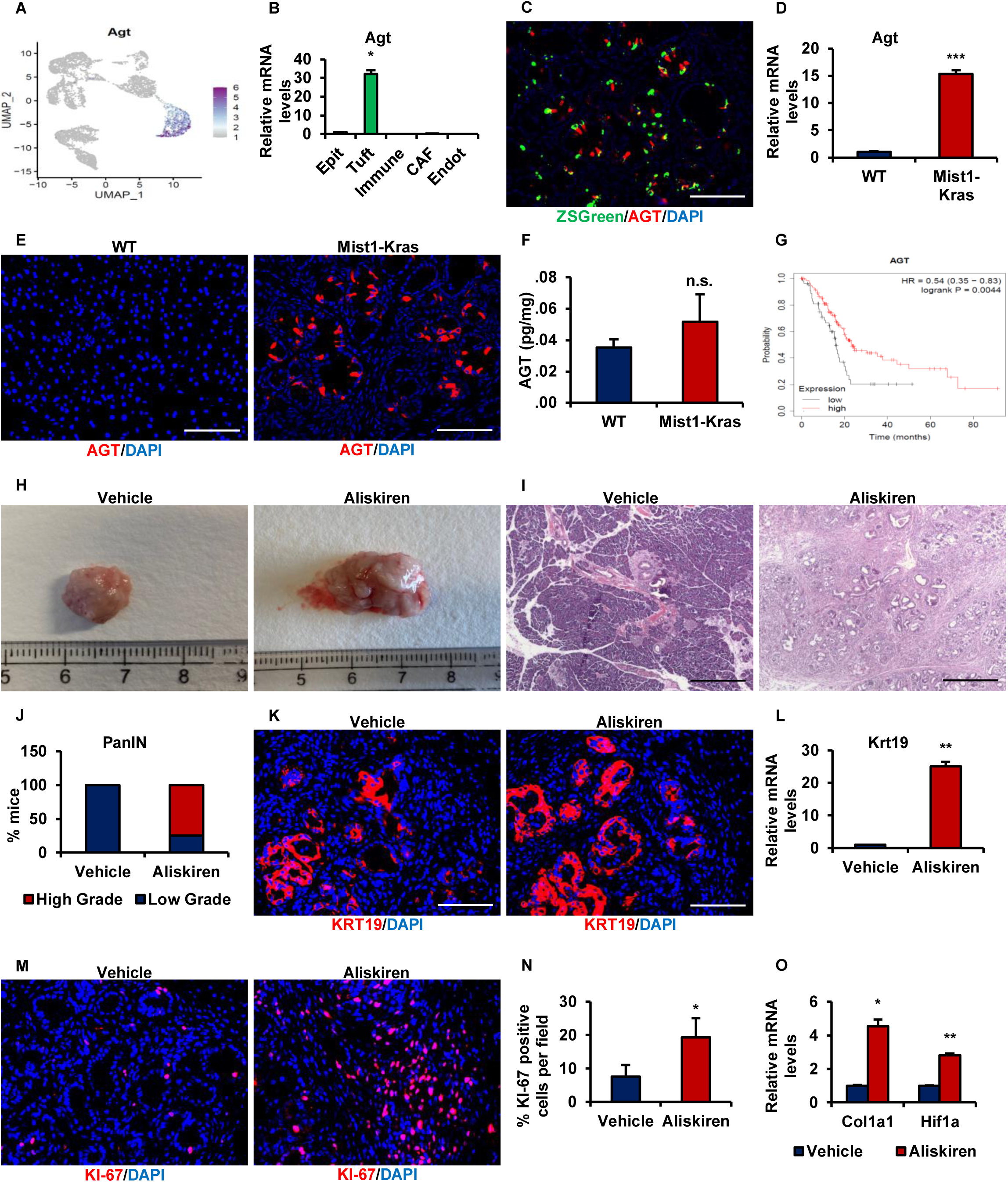
**(A)** UMAP showing expression of Agt in Dclk1^+^ cells. **(B)** qRT-PCR for *Agt* expression in epithelial cells (Epit), tuft cells, immune cells, CAFs and endothelial cells (Endot) isolated from Mist1-Kras-Dclk1-DTR-ZsGreen mice. **(C)** Immunofluorescence of pancreas from Mist1-Kras-Dclk1-DTR-ZsGreen mice for ZsGreen and AGT. **(D)** qRT-PCR for *Agt* expression in normal pancreas and pancreas from Mist1-Kras mice. **(E)** Immunofluorescence of normal pancreas and pancreas from Mist1-Kras mice for AGT. **(F)** ELISA of normal pancreas and pancreas from Mist1-Kras mice for AGT. **(G)** Kaplan-Meier plot of overall survival in PDAC patients based on low or high *Agt* expression. **(H)** Pancreas from Mist1-Kras mice treated with vehicle or aliskiren for 8 wks. **(I-J)** H&E of pancreas from Mist1-Kras mice treated with vehicle or aliskiren with related high or low grade PanIN quantification. **(K)** Immunofluorescence for KRT19 of pancreas from Mist1-Kras mice treated with vehicle or aliskiren. **(L)** qRT-PCR for *Krt19* expression in pancreas from Mist1-Kras mice treated with vehicle or aliskiren. **(M-N)** Immunofluorescence for KI-67 of pancreas from Mist1-Kras mice treated with vehicle or aliskiren and related quantification. **(O)** qRT-PCR for *Col1a1*, *Acta2*, *Cd31* and *Hif1a* expression in pancreas from Mist1-Kras mice treated with vehicle or aliskiren. Scale bars: 100 μm. Means ± SD. ∗: p ≤ 0.05; ∗∗: p ≤ 0.01; ∗∗∗: p ≤ 0.001.

We measured the expression levels of the *Agt* gene in several cell populations isolated from Mist1-Kras mice, and confirmed that tuft cells were the main source of this ligand in this model (Figure 7B). In agreement with this, immunofluorescence showed that AGT-producing cells displayed a tuft cell morphology as well as co-expression of ZsGreen (Figure 7C). The *Agt* gene was also included in the interaction network of SPIB *in vivo*, and its expression significantly correlated with SPIB protein activity and was increased in SPIB overexpressing pancreatic organoids (Figures S7A-B). Moreover, *Agt* expression was increased in pancreatic organoids grown in the presence of IL13 or ILC2, but reduced in Mist1-Kras mice treated with the anti-IL13 blocking antibody (Figures S7C-E).

*Agt* expression was increased in pancreas epithelium of Mist1-Kras mice compared to normal pancreas epithelium, supporting a potential role of this ligand in pancreatic neoplasia (Figures 7D-E). AGT protein levels as quantified by ELISA displayed an overall trend towards increased levels in Mist1-Kras mice, but also showed a high degree of variability, which might reflect the degradation of AGT into its downstream signaling products (Figure 7F). Higher levels of *Agt* expression correlated with improved overall survival in patients diagnosed with pancreatic ductal adenocarcinoma (PDAC), raising the possibility that angiotensinogen produced by Dclk1^+^ tuft cells might have a protective role against pancreatic tumor progression (Figure 7G).

To better evaluate the role of angiotensinogen in pancreatic tumors, we treated Mist1-Kras mice with the renin inhibitor aliskiren, giving that renin is the first enzyme in the degradation cascade of angiotensinogen into its downstream signaling peptides. The treatment was started at 2 weeks after KRAS activation and continued for a period of eight weeks. Mice treated with aliskiren displayed a marked accelerated pancreatic tumor progression, as revealed by significantly higher number of high-grade PanIN lesions and proliferating KI-67 cells compared to vehicle treated mice (Figures 7H-N). Aliskiren treatment also resulted in increased expression of markers of fibrosis (*Col1a1*, *Acta2*) as well as hypoxia (*Hif1a*), but decreased expression of the endothelial marker *Cd31* (Figure 7O).

We hypothesized that angiotensinogen could exert part of its effects on pancreatic tumor progression through angiotensin II. CAFs displayed the highest level of expression of angiotensin II receptors (*Agtr1*, *Agtr2*), indicative that they could be a main target of angiotensin II in pancreatic neoplasia (Figures S7F-I). In agreement with this possibility, we found that CAFs isolated from Mist1-Kras mice promptly responded to angiotensin II stimulation *in vitro*, as assessed by the marked increase in Tgf-β expression, a reported angiotensin II target gene (Figure S7J)^55,56^. On the other hand, this treatment did not affect the expression of genes for ECM proteins (*Col1a1*, *Acta2*) (Figures S7K-L).

Overall, our data show that angiotensinogen is a key signal provided by Dclk1^+^ tuft-like cells in early pancreatic neoplasia, where it plays important roles in restraining the progression of the disease to the more advanced stages.

## Discussion

We have used in our study a combination of a reporter mouse model, scRNA-seq and regulatory network analysis to decipher the identity of *Dclk1* expressing cells in normal pancreas and the molecular changes that occur in these cells during pancreatic tumorigenesis driven by mutant KRAS. In normal pancreas, *Dclk1* expression identifies epithelial cells localized in ducts, islets and acini. Dclk1^+^ cells in ducts exhibit features of terminally differentiated tuft-like cells, whose presence in the pancreas has been matter of debate, possibly due to the different sensibility of the methods used to study these rare cells^15,17^. Dclk1^+^ cells in the islets have features of neuroendocrine cells. Dclk1^+^ cells in the acini represent a subset of less differentiated acinar cells with peculiar neural and immune functions, which might serve as putative progenitor cells in response to pancreatic injury^8^. In contrast, in early pancreatic neoplasia single cell scRNA-seq analysis shows that the wide majority of Dclk1-ZsGreen^+^ cells can be classified into three clusters: acinar cells, Acinar-to-Ductal Metaplasia (ADM)-like cells and tuft-like cells. Dclk1^+^ acinar cells are primed to differentiate into ADM-like or tuft-like cells early during pancreatic tumor progression, as result of alternative KRAS-dependent intracellular pathways. Which of the two alternative differentiation pathways will be activated can be predicted by copy number variation (CNV) and genomic instability score (GI) of each acinar cell: cells with high KRAS activation and GI score will generate ADM-like cells; cells with low KRAS activation and GI score will generate tuft-like cells. The progression towards tuft-like cells is further orchestrated by an intercellular axis based on paracrine interactions involving IL13-secreting ILC2 cells and IL33-secreting cancer-associated fibroblasts (CAFs). Once generated, Dclk1^+^ tuft-like cells produce angiotensinogen that mediate their protective roles against the progression of pancreatic neoplasia.

Dclk1^+^ cells are markedly increased in early pancreatic neoplasia, but they decline in the more advanced stages of the disease. This increase is evident as early as 2 wks after KRAS activation and become more pronounced in the following weeks. *Kras*-mutant acinar cells sustain this expansion. This pool of *Kras*-mutant acinar cells presumably arise in part from the resident pool of Dclk1^+^ acinar cells in normal pancreas and in part from other Dclk1-negative acinar cells that begin to express *Dclk1* following KRAS activation. A key point of our study is that we provide evidence that *Dclk1* expression identifies five epithelial cell populations in early pancreatic neoplasia: ADM-like cells, tuft-like cells, acinar cells, progenitor cells, ductal cells. This finding therefore explains the reason of contrasting definitions that have attributed to Dclk1^+^ cells in pancreatic neoplasia^16,17^.

The acinar cluster is the largest cluster at 2 wks after KRAS activation, and includes acinar cells that express *Dclk1* following KRAS activation as well as normal Dclk1^+^ acinar cells. Although these two cell types co-cluster and it is not possible to discriminate them further based on gene expression, we found that these cells display intrinsic differences in terms of CNV and GI score, given that Kras mutant acinar cells show higher CNV and GI score with respect to Dclk1^+^ acinar cells, reflecting the different status of KRAS activation between them. This expanded pool of acinar cells resembles closely normal Dclk1^+^ acinar cells, therefore supporting the concept that Dclk1^+^ acinar cells could represent a subset of acinar cells with progenitor features in normal pancreas.

ADM-like cells and tuft-like cells represent the main Dclk1^+^ cell populations at 16 weeks after KRAS activation. ADM-like cells show low levels of *Dclk1* expression (ZsGreen^Low^), exhibit cancer stem cell markers (CD44^+^CD133^+^) and also the capacity to robustly generate organoids *in vitro*. The gene signature of Dclk1^+^ ADM-like cells is enriched in scRNA-seq profiles of human PDAC cells, supporting the concept that these cells have the potential to sustain pancreatic tumor progression. Tuft-like cells show high levels of expression of *Dclk1* (ZsGreen^Hi^) and other tuft cell markers, but lack of cancer stemness features, arguing against the possibility they could directly sustain pancreatic tumor growth. In line with this possibility, the signature of Dclk1^+^ tuft-like cells is not enriched in any scRNA-seq profiles of human PDAC cells we examined. This finding also supports the concept that tuft cells are most abundant in the early stages of pancreatic neoplasia but decline with the progression to PDAC.

ADM-like and tuft-like cells are both generated by *Kras*-mutant Dclk1^+^ acinar cells as the results of two alternative differentiation pathways, as revealed by pseudo-trajectory analysis. Each *Kras-*mutant acinar cell is primed to generate either a ADM-like cell or a tuft-like cell as early as 2 wks after KRAS activation. KRAS protein activity, CNV and GI score are critical to decide which differentiation pathway will be activated in each Dclk1^+^ acinar cell and therefore allow to predict this decision: cells showing high KRAS protein activity leading to higher CNV and GI score will differentiate into ADM-like cells; cells with low KRAS protein activity and lower CNV and GI score will rather generate tuft-like cells.

Dclk1^+^ tuft like-cells in pancreatic neoplasia display the activation of many well-known transcription factors reported to be responsible for tuft cell differentiation in other tissues (ASCL2, GFI1B, POU2F3, PROX1)^21,22,23,24^. However, we found of particular interest the ETS transcription factor SPIB, which it has not received much attention before, although consistently present in most of the reported tuft cell gene signatures^21,23,24,28^. In the intestine, SPIB has been shown to promote the differentiation of Lgr5 stem cells into microfold cells, a rare cell type involved in immune responses against non-self antigens^57,58^. However, not much is known about its roles in terms of tuft cell biology. We found SPIB of particular interest because it is among the first transcription factors to be significantly upregulated in *Kras*-mutant Dclk1^+^ acinar cells primed to differentiated in tuft-like cells, and also because it is expressed at high levels by Dclk1^+^ tuft-like cells, implying potential key roles of this transcription factor for both their differentiation and function. In agreement with this, we found that SPIB controls the expression of multiple tuft cell markers *in vivo* in pancreatic neoplasia as well as *in vitro* in pancreatic organoids (*Dclk1*, *Sox9*, *Cd24a*). In addition, SPIB regulates the expression of many genes involved in immunity and cytokine signaling (*Cxcl12*, *Il33*, *Cebpb*), indicative that this transcription factor might stimulate the activation of immune responses by Dclk1^+^ tuft-like in early pancreatic neoplasia, potentially with the aim to preserve tissue homeostasis from the neoplastic insult.

We found that the generation of Dclk1^+^ cells in pancreatic neoplasia depends not only on KRAS activation, but it is also highly dependent on exogenous signals from the tumor microenvironment. Given the link that has been described between tuft cells, ILC2 and IL4/IL13 in the intestine in context of helminthic infections^25,26,27^, we investigated whether this model could also apply to pancreatic neoplasia. Similarly to the intestine, we found that ILC2 sustain tuft cell generation in pancreatic neoplasia, through the secretion of IL13. In contrast to the intestine, where ILC2 are mainly activated by IL25 provided by tuft cells, we show that ILC2 are mainly activated by IL33 in pancreatic neoplasia, which is mainly provided by CAFs. A IL33-ILC2 axis has been identified in orthotopic mouse models of pancreatic tumors and reported to play protective roles against tumor growth by stimulating CD8 T cell activation^33^. Our data therefore support the notion that Dclk1^+^ tuft-like cells, as well as CAFs, participate in this intercellular axis existing in pancreatic neoplasia, where they could reinforce the protective response against tumor progression. Recent studies have pointed out roles for the tuft cells-ILC2 axis in promoting gastric tumor formation based on lineage depletion of Dclk1^+^ cells or cytokine blocking experiments^48^.

We show here that Dclk1^+^ cells represent a heterogeneous population of cells including tuft cells as well as other epithelial cell types that could promote tumor progression, such as ADM-like cells, indicative that an indiscriminate depletion of Dclk1^+^ cells is not well-specific to address the roles of tuft-like cells in neoplasia. In addition, the effects of blocking cytokines involved in the tuft cell-ILC2 axis could be also dependent on side effects on other immune cell types, as also reported in previous studies^48^. Our findings pointing out to the protective roles of the tuft cell-ILC2 axis against pancreatic neoplasia together with the well-established roles of this axis against helminthic infection of the intestine lets speculate that some form of infections might be protective against some types of tumors, especially of the gastrointestinal tract, and could partially explain the increased tumor incidence over the centuries.

Previous studies have indicated that tuft cells contribute to inhibit pancreatic tumorigenesis by producing Prostaglandin D2 (PGD2), a lipid mediator synthesized by the enzyme HPGDS. PGD2 would suppress the inflammatory response mediated by macrophages as well as pancreatic stellate cell activation, and by this would inhibit pancreatic tumor progression^20^. Our Dclk1^+^ tuft-like cell signature includes several genes encoding for enzymes involved in the production of lipid mediators of the prostaglandin and leukotriene families, including *Alox5ap*, *Hpgds* and *Ptgs1*, therefore confirming a role for these signals in pancreatic neoplasia. However, we decided to focus our attention on angiotensinogen because it was among the only few ligands produced at high levels by Dclk1^+^ tuft-like cells in pancreatic neoplasia. Furthermore, we found this ligand to be consistently reported in tuft cell signatures of other organs, which supported the possibility of its potential key roles for tuft cell function^21,23^. In addition, Dclk1^+^ tuft-like cells are the main source of angiotensinogen in pancreatic neoplasia, therefore making this ligand an ideal target to assess the role of these cells in this disease. We found angiotensinogen to be a key inhibitory signal provided by Dclk1^+^ tuft-like cells against the progression of pancreatic neoplasia. To demonstrate this, we employed aliskiren, a renin inhibitor that blocks the degradation of angiotensinogen into its downstream signaling peptides. Remarkably, this treatment results in markedly accelerated pancreatic tumor progression, with significant higher numbers of high-grade PanIN lesions and proliferating KI-67 cells as well as increased fibrosis and hypoxia. To further support our finding, Kaplan-Meier survival analysis shows that PDAC patients expressing higher levels of angiotensinogen have a significantly better overall survival with respect to the other patients. The role of angiotensinogen itself has not been well investigated in cancer, although studies have reported roles for angiotensin II in various established tumor models ^50,51,52,53,54^. These studies have largely focused on established tumors and on the benefits of blocking angiotensin II either by ACE inhibitors or sartans in order to reduce ECM stiffness and improve chemotherapy delivery to the tumor, therefore improving the efficacy of therapies. However, studies have not addressed the roles of angiotensin II during tumor progression, and more specific at the early stages neoplasia, which therefore remains to be defined. In light of these studies, we speculated that angiotensinogen could exert its protective effects in early pancreatic neoplasia at least in part through angiotensin II, where CAFs would represent the main cellular target. To support this possibility, we found that CAFs promptly respond to angiotensin II stimulation by upregulating *Tgfβ* expression, which is known to play pleiotropic roles in tumor progression, with tumor suppressive roles at the early stages and tumor promoting roles at the later stages^59^. However, it plausible that other downstream signaling peptides originated by angiotensinogen degradation could mediate its effects in pancreatic neoplasia, which remain to be identified and investigated more comprehensively.

In conclusion, we describe here a novel tuft cell-ILC2 axis that plays a key role to inhibit pancreatic tumor progression, therefore supporting the emerging concepts of the protective roles of tuft cells and ILC2 in pancreatic cancer^20,33^. The tuft cell-ILC2 axis is possibly part of a broader protective response of the pancreas against neoplastic transformation, which also involves other immune cells, like macrophages, dendritic cells, CD8 T cells as well as CAFs^20,33^. In this scenario, tuft cells contribute at least in two ways: they sustain ILC2 activation together with CAFs by providing IL33; they produce mediators that directly suppress the progression of pancreatic tumors, including angiotensinogen. The involvement of angiotensinogen against pancreatic neoplasia has broader clinical implications, as it also suggests that some drugs targeting the RAAS pathway and widely used as anti-hypertensive should be better evaluated retrospectively in terms of potential susceptibility to some form of cancer, including PDAC. Collectively, our study provides insights for the development of novel therapeutic strategies that could potentially exploit the tuft cell-ILC2 axis for the treatment of pancreatic tumors.

## Experimental Procedures

### Animal Models

Dclk1-DTR-ZsGreen mice were generated in our laboratory. Briefly, the cassette DTR-2A-ZsGreen-pA-FrtNeoFrt was inserted into a pL451 plasmid and electroporeted into SW105 cells carrying the Dclk1-BAC vector (clone RP23-283D6) in order to obtain recombination downstream the ATG of exon 2 of the mouse *Dclk1* gene. Recombinant BAC vectors were isolated, linearized and microinjected into the pronucleus of fertilized CBA × C57BL/6J oocytes at the Columbia University Transgenic Animal Core facility and backcrossed into C57BL/6J mice. Mist1-CreERT2, Rosa26-tdTomato and wild-type mice were obtained from the Jackson Laboratory. Pdx1-Cre, lsl-KRAS^G12D^ and lsl-p53^R172H^ were provided by Dr. Kenneth P. Olive at Columbia University. All mice were housed in a pathogen-free facility under regular chow diet. All mice were treated starting at an age of 8 wks and sacrificed at different time points depending on the study and as described in the main text. Mice of both sexes were used in the study. All mouse studies were performed in accordance with the guidelines approved by the Columbia University Institutional Animal Care and Use Committee (IACUC).

### Mouse Treatments

To induce transgene expression in appropriate mouse lines, mice were induced once with tamoxifen (Sigma) at a single dose of 4 mg dissolved in corn oil by oral gavage. For IL13 blocking experiment, IL13 blocking antibody (R&D) or isotype control (R&D) were dissolved in DPBS (Thermo Fisher) and administered intraperitoneally every other day at 1 mg/kg for a period of 8 wks. To inhibit renin, aliskiren (Sellckchem) was dissolved in DPBS and administered at 25 mg/kg/day via osmotic minipumps model 2004 (Alzet). Mice were anesthetized using isoflurane, skin incisions were subsequently made on the back of the animals and osmotic minipumps were implanted into a subcutaneous pocket in order to deliver either aliskiren or DPBS continuously for a period of 8 wks and replaced after 4 wks.

### Cell isolation from murine pancreas

Mice were sacrificed according to IACUC protocol. Pancreas was removed and transferred on ice-cold HBSS (Thermo Fisher). Pancreas was then dissociated with a scalpel on ice until getting small pieces of tissue of approximately 1-2 mm. Pieced were then collected and digested in 1 mg/ml Collagenase P (Sigma) at 37°C for 20 minutes. After washing, samples were further digested in Accutase (Thermo Fisher) at 37°C for 5 minutes. After washing, samples were filtered with 40 µm cell strainers (BD) on ice. Red blood cell lysis was then performed by adding ACK lysing buffer (Thermo Fisher) to the cell suspensions and incubating samples on ice for 5 minutes. After washing, cells were manually counted and further processed for downstream applications.

### Flow cytometry and sorting

Single cell suspensions were resuspended in flow cytometry buffer containing DPBS supplemented with 0.5% BSA Fraction V Molecular Biology Grade (Gemini). Antibodies were added at the appropriate dilutions and incubated on ice in the dark for 30 minutes. After washing, samples were resuspended in flow cytometry buffer and vital staining was performed by adding DAPI (BD). For intracellular staining of IL13 and IL4, samples were fixed and permeabilized with FIX & PERM Cell Fixation and Cell Permeabilization Kit (Thermo Fisher) according to manufacturer’s instructions. Samples were analyzed on BD Fortessa or sorted with BD FacsAria II. Data were analyzed with FlowJo.

### Organoid and non-adherent sphere cultures

For organoids cultures, single cell suspensions from mouse pancreas were mixed with Matrigel-GFR (Corning) and seeded at 10000 cells/100 µl Matrigel/well in 24-well plates on ice. Plates were then incubated at 37°C for 30 minutes to allow Matrigel solidification. Subsequently, wells were overlaid with 500 µl of culture medium. Cultures were maintained in humidified atmosphere with 5% CO2 at 37°C for 7 days with medium changes every other day. Cells were grown in Advanced DMEM/F12 (Thermo Fisher) supplemented with Glutamax (Thermo Fisher), HEPES (Thermo Fisher), B27 (Thermo Fisher), N2 (Thermo Fisher), Antibiotic/Antimicotic (Thermo Fisher), 20 ng/ml EGF (Peprotech), 20 ng/ml bFGF (Peprotech). For the initial plating, 10 µM ROCK inhibitor (Sigma) was also added to the medium. For IL13 stimulation, 20 ng/ml IL13 (Peprotech) was added to the medium every other day for 7 days. For ILC2 co-cultures, 250-500 ILC2 isolated from Mist1-Kras mice with 20 ng/ml IL2 (Peprotech) and 20 ng/ml IL7 (Peprotech) were added to the growing medium at the day of the plating in 96-well plates. For experiments involving viral infections, single cell suspensions in Matrigel were added with Spib overexpressing lentivirus (VectorBuilder) or Adeno-CMVCre (University of Iowa) at 500 PFU/cell and subsequently grown as described above. To select for lentiviral transfected organoids 2 µg/ml puromycin (Sigma) was added to the medium.

For non-adherent sphere cultures, single cell suspensions from mouse pancreas were seeded at 20000/well in 24-well plates coated with poly-HEMA (Sigma). Cells were grown for 7 days in same culture medium as for organoids with medium changes every other day.

Bright field and fluorescence images of organoids and non-adherent spheres were acquired with an Eclipse TU2000-U microscope (Nikon).

### Cell cultures

ILC2 (Lin^-^CD45^+^CD127^+^CD90^+^KLRG1^+^) were sorted from pancreas of Mist1-Kras mice and grown in DMEM high glucose, Glutamax supplement and pyruvate (Thermo Fisher) supplemented with HEPES, 10% FBS (Thermo Fisher), Antibiotic/Antimycotic, 20 ng/ml IL2, 20 ng/ml IL7. For IL33 stimulation, 20 ng/ml IL33 (Peprotech) was added to the medium.

CAFs (PDGFRA^+^) were sorted from pancreas of Mist1-Kras mice and grown in in DMEM high glucose, Glutamax supplement and pyruvate (Thermo Fisher) supplemented with HEPES, 10% FBS (Thermo Fisher), Antibiotic/Antimycotic. For angiotensin 2 stimulation, 1 µM Ang II (Sigma) was added to the medium for 48 h.

### Single-cell gene RNA Sequencing Analysis

Single-cell RNA sequencing was performed on sorted cells from mouse pancreas. Data were analyzed using the 10X Cell Ranger software (10x Genomics). Samples were combined using the Cell Ranger aggregate function.

Single cell data were filtered for low quality cells. Cells with less than 1000 UMI and/or a high fraction of mitochondrial counts (>30%) were removed. Single cell gene expression profiles were normalized to count per millions (CPM), transformed to gene expression signatures by z-score transformation and used to generate meta-cell profiles that were used only as input for ARACNe^60^. Specifically, z-score expression signatures were used to perform a K-nearest neighbors (KNN) analysis and identify for each individual cell the closest 10 cells. The KNN analysis was performed in the distance matrix generated by reciprocal enrichment analysis of the 200 most differentially expressed genes (100 most overexpressed and 100 most downregulated) as implemented in the viperSimilarity function of the VIPER package. A metacell profile was then generated for each cell by integrating the raw counts of the 10 closest cells (KNN approach) in the viperSimilarity space. To avoid potential bias due to different microenviroment Dclk1^+^ normal cells were not used to generate metacells. Metacell profiles were normalized and used only as input for ARACNe to generate the regulatory network.

ARACNe was run with 100 bootstrap iterations and default parameters using 1824 transcription factors (genes annotated in gene ontology molecular function database, as GO:0003700, “transcription factor activity”, or as GO:0003677, “DNA binding”, and GO:0030528, “transcription regulator activity”, or as GO:00034677 and GO: 0045449, “regulation of transcription”) and 3477 signaling pathway-related genes (annotated in GO biological process database as GO:0007165 “signal transduction” and in GO cellular component database as GO:0005622, “intracellular”, or GO:0005886, “plasma membrane”). ARACNe inferred a regulatory network (Dclk1 network) of 3770 protein regulons.

### VIPER analysis

The Dclk1 network was used for VIPER analysis and to infer protein activity profiles from single cell gene expression profiles. Specifically, normalized single-cell gene expression profiles were first transformed in gene expression signatures by z-score transformation then converted to protein activity profiles by VIPER using the ARACNe networks. To identify cell types, we performed unsupervised cluster analysis based on VIPER-inferred protein activity profiles and on gene expression profiles using the Seurat package. For cluster analysis we used the Louvain algorithm following the clustering procedure implemented in the Seurat package.

The optimal number of clusters was estimated by varying the resolution parameter (from 0.1 to 1 with intervals of 0.05) of the “FindCluster” function and cluster solutions were evaluated by silhouette analysis. Since protein activity clustering outperformed gene expression clustering based on silhouette analysis we determined cell types based on the optimal number of clusters estimated by silhouette analysis on the protein activity profiles. Gene expression analysis including differential gene expression analysis was performed using Seurat (V4.3).

Master Regulator Analyses (MRA) was performed using the msviper function of the VIPER package by applying the Dclk1 network on the gene expression signature computed by comparing the gene expression centroids of conditions.

Gene set enrichment analyses were performed using the either one-tailed of two-tailed test with 1000 permutations. aREA tests were performed using the aREA function implemented in the VIPER package. Genomic Instability Analysis was performed using the genomic instability package available on Bioconductor (https://www.bioconductor.org/packages/release/bioc/html/genomicInstability.html)^45^.

### RNA-seq

Organoids were processed with the RNeasy Plus Mini kit (Qiagen) according to the manufactureŕs instructions. RNA quantification and quality assessment were evaluated by Bioanalyzer (Agilent): all samples used in the experiment had a RIN of 10. poly-A pull-down was used to enrich mRNAs from total RNA samples and then library was constructed using Illumina TruSeq chemistry. Libraries were then sequenced using Illumina NovaSeq 6000. Samples were multiplexed in each lane, yielding targeted number of paired-end 100bp reads for each sample. RTA (Illumina) was used for base calling, and bcl2fastq2 (version 2.19) for converting BCL to fastq format, coupled with adaptor trimming. A pseudoalignment was performed to a kallisto index created from transcriptomes (Mouse: GRCm38) using kallisto (0.44.0). Data were further analyzed with Partek for differential gene expression among samples.

### qRT-PCR

Whole tissues were processed with RNeasy plus mini kit (Qiagen) according to the manufactureŕs instructions. Sorted cells were processed with the RNeasy Plus Micro kit (Qiagen) according to the manufactureŕs instructions. RNA quantification was performed with Nanodrop (Thermo Fisher). RNA quantification was performed with Nanodrop (Thermo Fisher). RNA (500 ng to 2 µg) was further processed to synthesize cDNA with SuperScript III Reverse Transcriptase (Thermo Fisher) according to the manufactureŕs instructions. Real time PCR was run using PowerUp SYBR Green master mix (Thermo Fisher) according to the manufactureŕs instructions. Quantitative PCR was performed in triplicate on an Applied Biosystems QuantStudio 3 machine. Relative gene expressions were normalized to Rpl19 or Actb.

### Survival analysis

Kaplan-Meier overall survival analysis of PDAC patients was evaluated using Kaplan-Meier Plotter (https://kmplot.com)

### Histology and Immunofluorescence

Dissected murine pancreas was fixed in 4% paraformaldehyde (PFA), embedded in OCT and snap frozen in liquid nitrogen. Frozen blocks were then sectioned on slides at 5 µm thickness.

For immunofluorescence, slides were washed in DPBS, blocked and permeabilized with 1% BSA/1% Triton X-100 (Fisher) in DPBS at room temperature (RT) for 1 h. Primary antibodies were added in 1% BSA in DPBS and incubated at 4°C overnight. After washing, secondary antibodies conjugated with fluorophores (Thermo Fisher) were added in 1% BSA in DPBS and incubated at room temperature in the dark for 1 hour. After washing, were counterstained and mounted with Vectashield antifade DAPI-containing mounting medium (Vector Laboratories). Fluorescence images were acquired with an Eclipse TE2000-U microscope (Nikon). For quantification studies, three random fields were acquired and staining was then quantified. Images were processed and analyzed with ImageJ.

For histology studies, slides were washed and stained for hematoxylin and eosin. Histological scoring of PanIN lesions was based on cellular atypia and proliferation according to published criteria and performed on samples whose identities were not revealed until the end of the evaluation. Low-grade PanIN lesions were defined as PanIN-1 and PanIN-2, high-grade PanIN lesions as PanIN-3.

### ELISA

Whole pancreatic tissues were processed with RIPA buffer (Thermo Fisher). Protein concentration was determined with Pierce BCA protein assay kit (Thermo Fisher) on a 96-well plate format and absorbance measured at 560 nm with Fisher Scientific Multiskan FC plate reader.

Murine Angiotensinogen (IBL-America) and IL33 (R&D) ELISA were performed according to manufacturer’s instructions. Absorbance was measured at 405-nm absorbance with Fisher Scientific Multiskan FC plate reader. Samples were run in triplicates.

### Statistical Analysis

Statistical measures include mean values and standard deviation error. Statistical significance was evaluated by Student’s t-test and defined for P values < 0.05 between groups.

## Acknowledgments

This work was supported by grants from the NIH and DOD including R35CA210088, UO1DK103155, RO1DK128195, W81XWH-21-1-0901, and the Pancreatic Cancer Action Network (PanCan/AACR) grant to T.C.W.; as well as an NCI Outstanding Investigator Award (R35 CA197745) and two NIH Shared Instrumentation Grants (S10 OD012351 and S1 0OD021764) to A.C.. This research was funded in part through the NIH/NCI Cancer Center Support Grant P30CA013696 and used the resources of the Herbert Irving Comprehensive Cancer Center Flow Cytometry Shared Resources, Molecular Pathology/MPSR, Genomics and High Throughput Screening, as well as the Genetically Modified Mouse Model Shared Resource (GMMMSR). This research was supported in part by the Columbia University Digestive and Liver Disease Research Center (CU-DLDRC) grant 1P30DK132710 and its Bio-Imaging, Organoid/Cell Culture, and Bioinformatics/Single-Cell Analysis Cores. R.T. was supported by Princess Takamatsu Cancer Research Fund, Takeda Science Foundation, Astellas Foundation for Research on Metabolic Disorders, and AMED under Grant Number JP22cm0106284. H.N. was supported by Deutsche Forschungsgemeinschaft Grant NI 1810/1-1. We are grateful for the support we received by Columbia University shared resources and we would like to thank Sun Dajiang “Kevin” (Molecular Pathology/MPSR), Kissner, Michael (Columbia Stem Cell Initiative Flow Cytometry core facility), Erin Bush (Genomics and High Throughput Screening), and Lin Chyuan-Sheng “Victor” (Genetically Modified Mouse Modeling), for their incredible expertise and help in this project. We thank Nicoletta Barolini (Columbia University) for designing the graphical models.

## Author Contributions

Conceptualization: G.V., P.L., T.C.W.; Methodology: G.V., P.L., R.T., Y.H., Investigation: G.V., P.L., R.T., F.W., T.R., Z.J., M.S., M.M., H.N., N.F., E.M., O.C.; Formal Analysis: G.V., P.L., A.V., S.T., A.C.I.; Writing: G.V., P.L., T.C.W.; Supervision: A.C., T.C.W.

## Declaration of Interests

P.L. is director of Single-Cell Systems Biology at DarwinHealth Inc., a company that has licensed some of the algorithms used in this article from Columbia University. A.C. is founder, equity holder, and consultant of DarwinHealth Inc. Columbia University is also an equity holder in DarwinHealth Inc.

---

**Figure S1.**
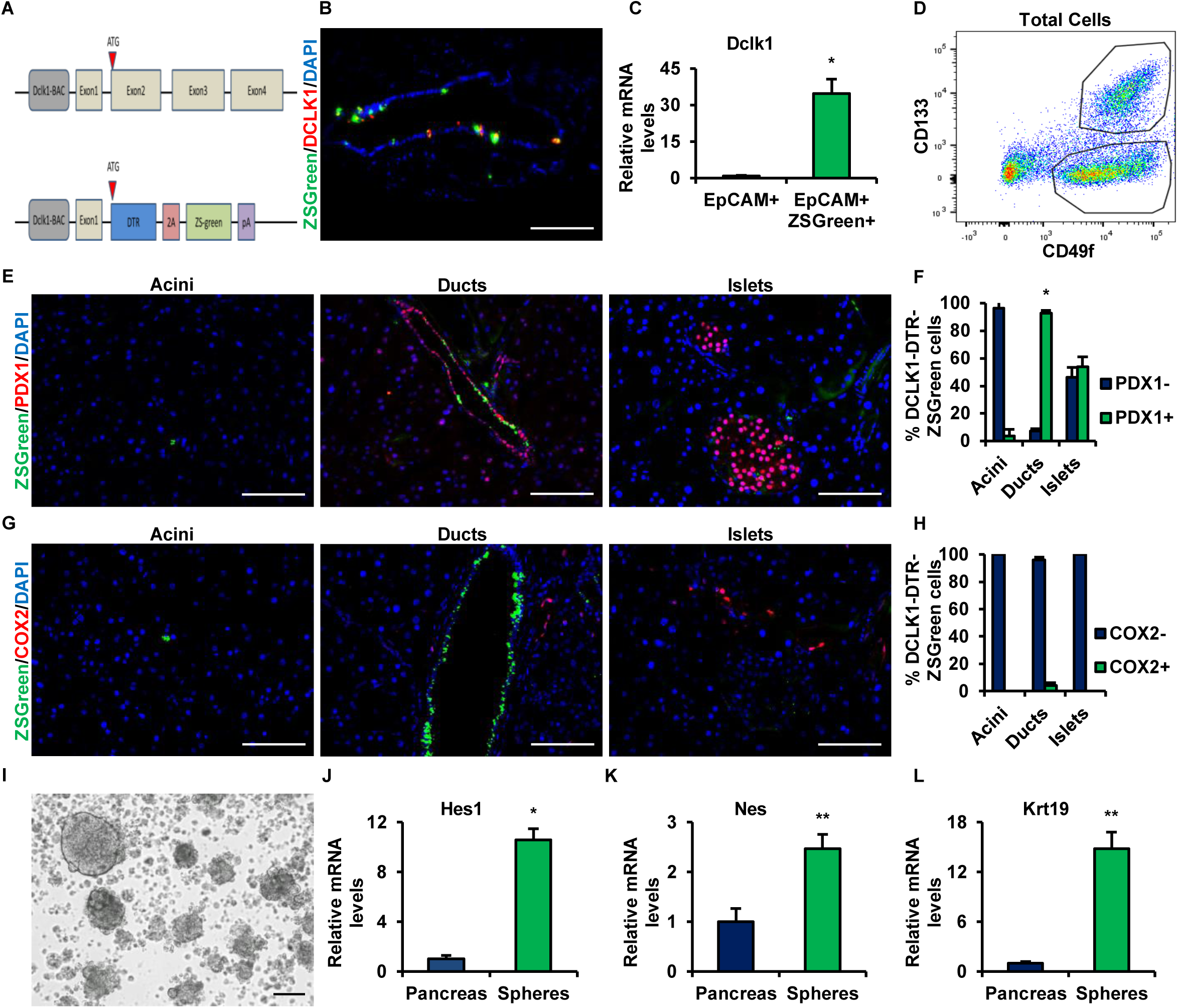
**(A)** Cartoon showing the construct to generate Dclk1-DTR-ZsGreen mice. **(B)** Immunofluorescence of normal pancreas of Dclk1-DTR-ZsGreen mice for DCLK1. **(C)** qRT-PCR for expression of *Dclk1* in EpCAM^+^ and EpCAM^+^ZsGreen^+^ cells isolated from normal pancreas of Dclk1-DTR-ZsGreen mice. **(D)** Flow cytometry analysis of single cells isolated from normal pancreas of Dclk1-DTR-ZsGreen mice for CD133 and CD49f. **(E-H)** Immunofluorescence of normal pancreas of Dclk1-DTR-ZsGreen mice for PDX1 and COX2 and related quantifications. **(I)** Non-adherent spheres grown from single cells isolated from normal pancreas of Dclk1-DTR-ZsGreen mice. **(J-L)** qRT-PCR for expression of *Hes1*, *Nes* and *Krt19* in non-adherent spheres. Scale bars: 100 μm. Means ± SD. ∗: p ≤ 0.05; ∗∗: p ≤ 0.01; ∗∗∗: p ≤ 0.001.

**Figure S2.**
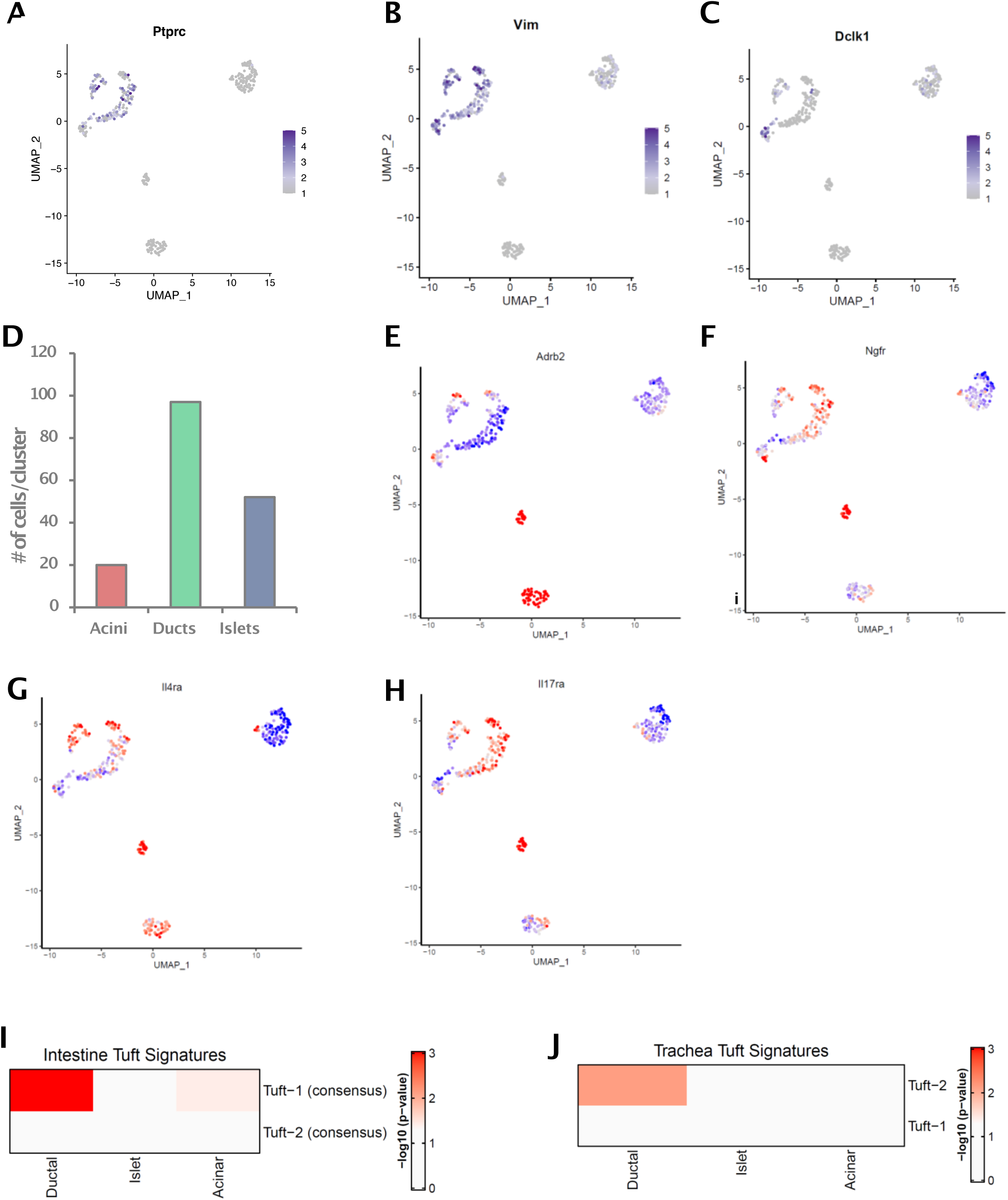
**(A-C)** UMAP plots showing the expression of *Ptprc*, *Vim* and *Dclk1* in Dclk1^+^ normal cells. **(D)** Barplot showing the number of cells in acinar, ducts and islet clusters. **(E-H)** UMAP plots showing Viper inferred protein activity of *Adrb2*, *Ngfr*, *Il4ra* and *Il17ra* in Dclk1^+^ normal cells. **(I-J)** Heatmaps showing the enrichment for Tuft1 and Tuft2 gene expression signatures derived from intestine and trachea in the gene expression signature Ductal, Islet and Acinar clusters identified in Dclk1^+^ normal cells. P-values were estimated by 1-tailed GSEA test with 1000 permutations.

**Figure S3.**
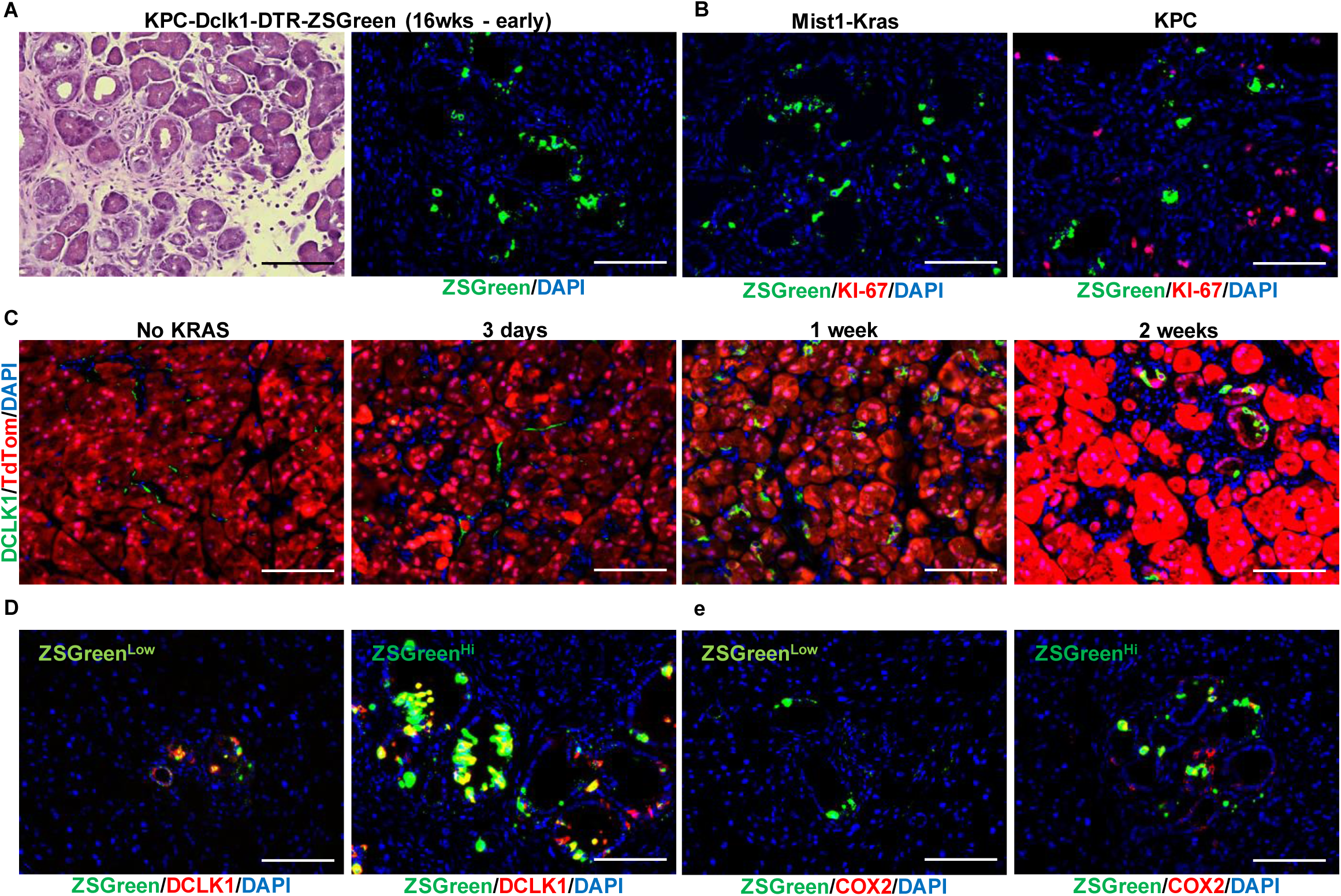
**(A)** Hematoxylin and Eosin staining (H&E) of pancreas from KPC-Dclk1-DTR-ZsGreen mice (area with early neoplasia) at 16 wks. **(B)** Immunofluorescence of pancreas from Mist1-Kras-Dclk1-DTR-ZsGreen mice and KPC-Dclk1-DTR-ZsGreen mice for KI-67. **(C)** Immunofluorescence of pancreas of Mist1-Kras-TdTomato mice not tamoxifen induced (No KRAS) and 4 wks, 8 wks and 16 wks after tamoxifen administration for DCLK1 and TdTom. **(D-E)** Immunofluorescence of pancreas of Mist1-Kras-Dclk1-ZsGreen mice for DCLK1 and COX2 in ZsGreen^low^ and ZsGreen^hi^ cells. Scale bars: 100 μm.

**Figure S4.**
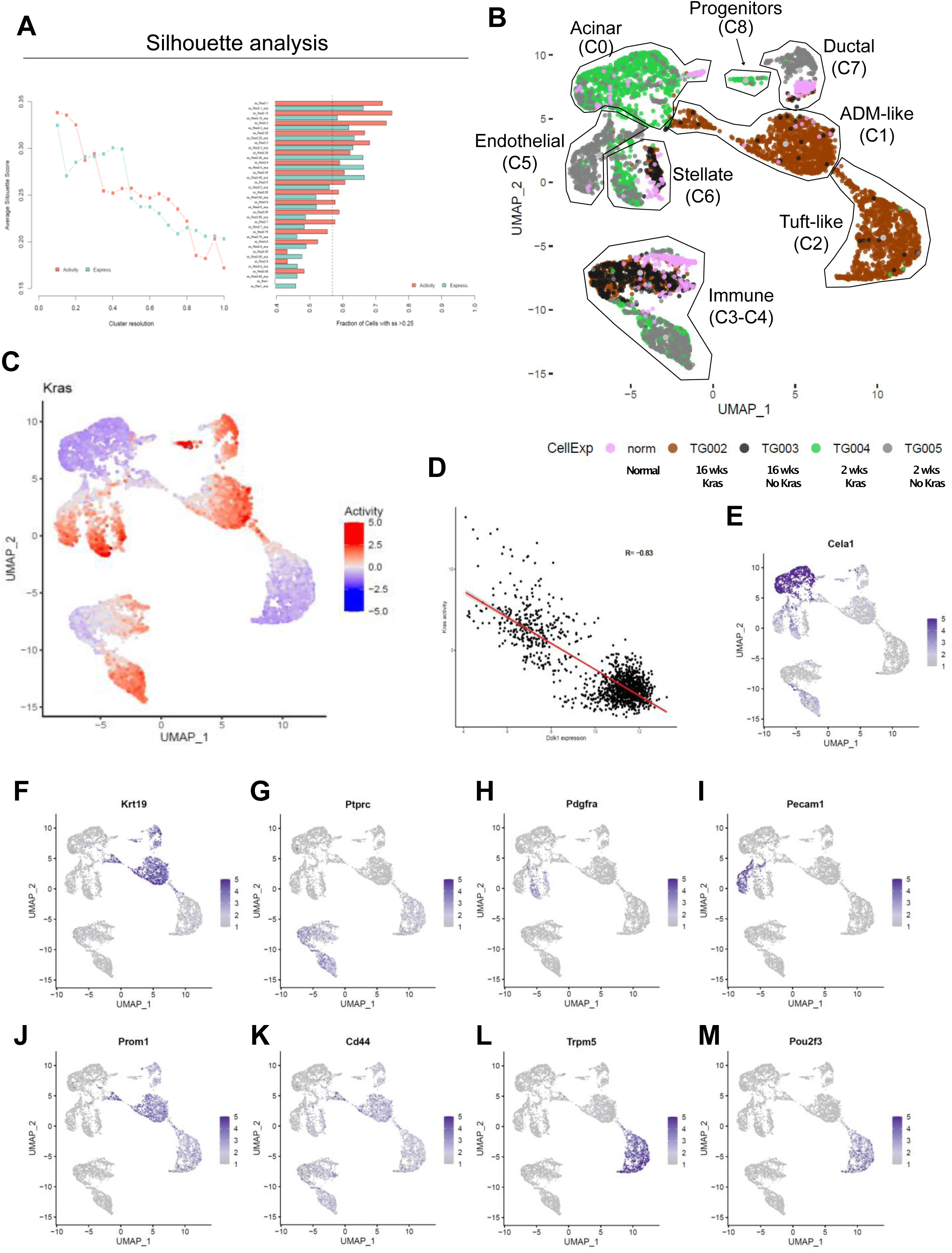
**(A)** Silhouette analysis for clustering optimization. Left, scatterplot showing the average silhouette scores computed by varying the resolution parameter (from 0.1 to 1 at intervals of 0.05) of the “FindCluster” function implemented in the Seurat package. Red line represents cluster solutions computed on VIPER inferred protein activity profiles. Cyan line represents cluster solutions computed on gene expression profiles. Right, Barplot showing the fraction of cells with a silhouette score > 0.25 for each solution. **(B)** UMAP based on VIPER activity showing the clusters/cell types identified across all single-cell experiments performed in this study. Cells derived from different experiment were annotated with different colors. **(C)** UMAP showing VIPER-inferred differential protein activity of KRAS. **(D)** Scatterplot showing the inverse correlation between *Dclk1* expression and VIPER-inferred KRAS protein activity. **(E-M)** UMAP plots showing the expression of cell type marker genes.

**Figure S5.**
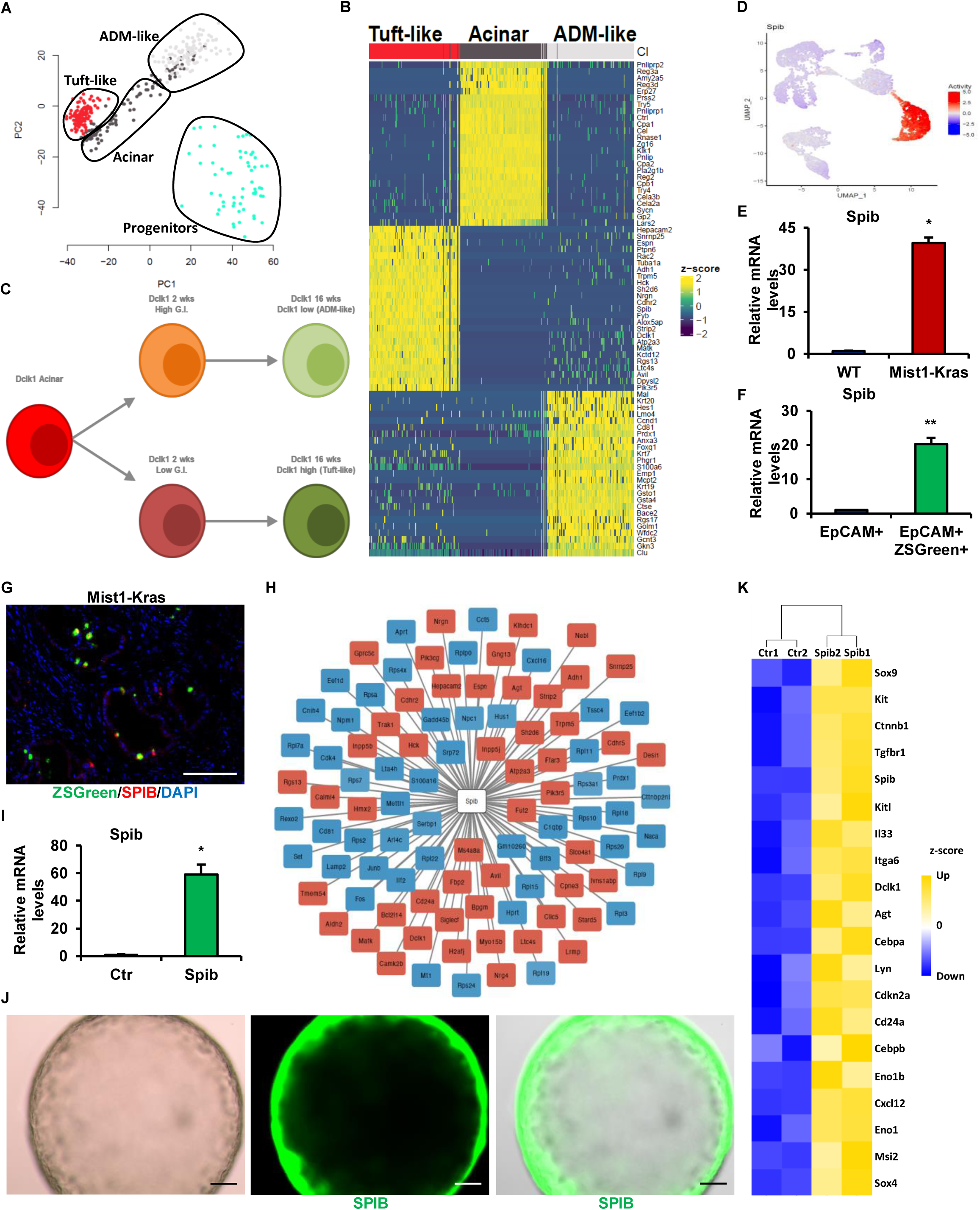
**(A)** PCA plot showing Tuft-like, Acinar, ADM-like and Progenitor cells. **(B)** Heatmap showing differential expressed genes of Tuft-like, Acinar and ADM-like cells. Columns represent cells and are sorted based on the pseudo trajectory. Expression is expressed as z-score values. **(C)** Cartoon representing results of GIS and pseudo trajectory analysis (Fig.5A, 5G) with the subset of acinar cells primed towards tuft-like cells showing lower GIS and acinar cells primed towards ADM-like cells showing higher GIS. **(D)** UMAP plot showing the expression of *Spib* in Dclk1^+^ cells. **(E)** qRT-PCR showing *Spib* expression in normal pancreas (WT) and early pancreatic neoplasia (Mist1-Kras). **(F)** qRT-PCR showing *Spib* expression in EpCAM^+^ and EpCAM+ZsGreen^+^ cells in Mist1-Kras mice. **(G)** Immunofluorescence staining of early pancreatic neoplasia (Mist1-Kras) for SPIB. **(H)** Gene network analysis of *Spib* in Dclk1^+^ cells inferred from scRNA-seq data. **(I)** qRT-PCR showing *Spib* expression in organoids generated from single cells isolated from pancreas of Kras mutant mice and transfected with control (Ctr) or *Spib*-expressing lentivirus (Spib). **(J)** Immunofluorescence evaluation of eGFP expression in *Spib*-overexpressing organoids. **(K)** Heatmap showing top genes differentially expressed in *Spib*-overexpressing organoids with respect to control organoids. Expression is represented by z-score. Scale bars: 100 μm. Means ± SD. ∗: p ≤ 0.05; ∗∗: p ≤ 0.01; ∗∗∗: p ≤ 0.001.

**Figure S6.**
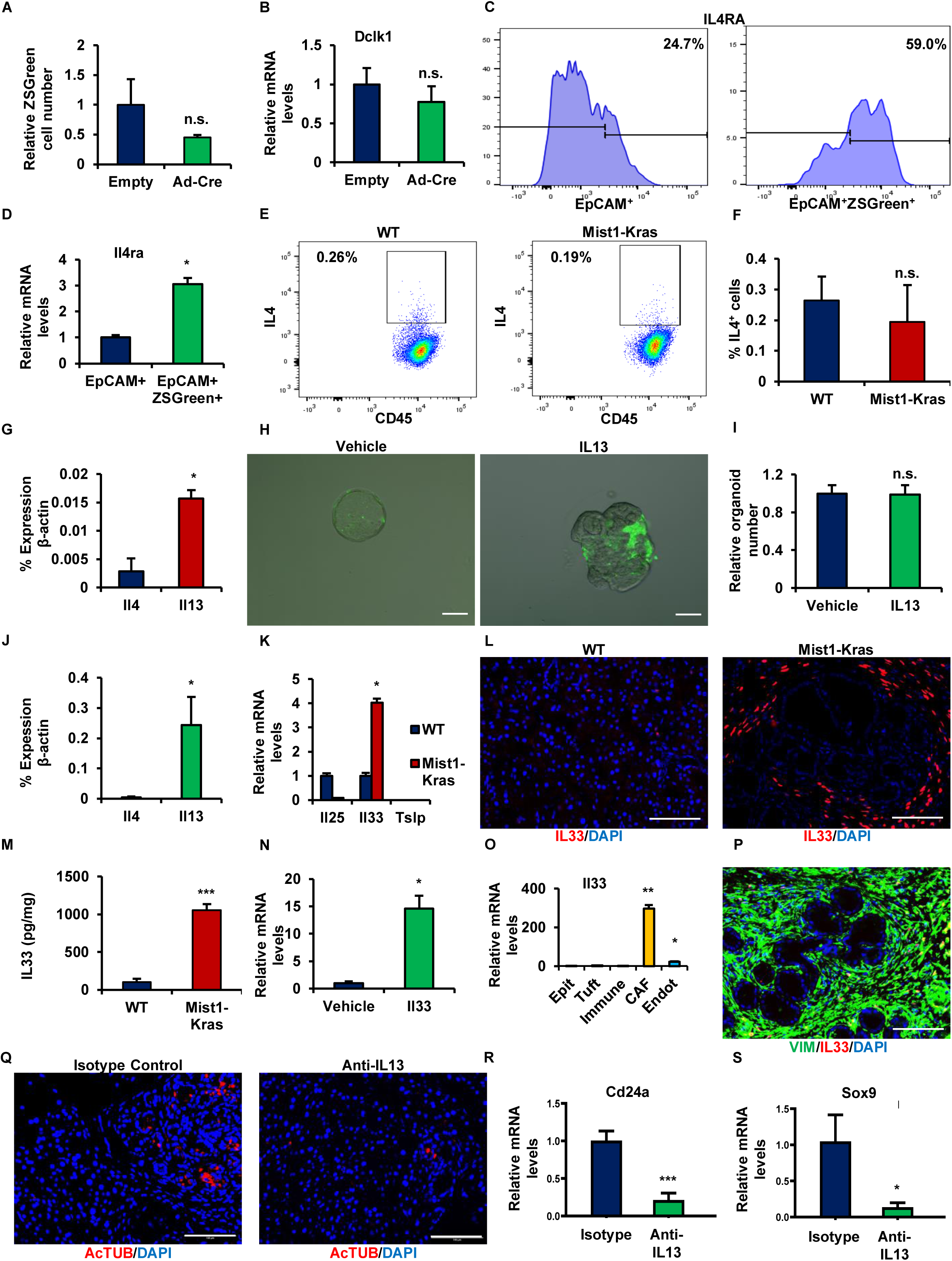
**(A)** ZsGreen expression in organoids grown from single cells isolated from pancreas of Kras-Dclk1-DTR-ZsGreen mice in the presence of empty adenovirus or Adeno-Cre adenovirus (Ad-Cre). **(B)** qRT-PCR for *Dclk1* expression in organoids grown in the presence of empty adenovirus or Ad-Cre. **(C-D)** Flow cytometry analysis for IL4RA in EpCAM^+^ or EpCAM^+^ZsGreen^+^ cells isolated from pancreas of Mist1-Kras-Dclk1-DTR-ZsGreen mice and related quantification. **(E-F)** Flow cytometry analysis for IL4 expression in immune cells (CD45) isolated from normal pancreas and pancreas from Mist1-Kras mice and related quantification. **(G)** qRT-PCR for *Il4* and *Il13* expression in pancreas of Mist1-Kras mice. **(H)** ZsGreen expression in organoids grown from single cells isolated from pancreas of Kras-Dclk1-DTR-ZsGreen mice in the presence of vehicle or IL13. **(I)** Relative number or organoids grown in the presence of vehicle or IL13. **(J)** qRT-PCR for *Il4* and *Il13* expression in ILC2 cells isolated from pancreas of Mist1-Kras mice. **(K)** qRT-PCR for *Il25*, *Il33* and *Tslp* expression in normal pancreas and pancreas from Mist1-Kras mice. **(L)** Immunofluorescence of normal pancreas and pancreas from Mist1-Kras mice for IL33. **(M)** ELISA of normal pancreas and pancreas from Mist1-Kras mice for IL33. **(N)** qRT-PCR for *Il13* expression of ILC2 isolated from Mist1-Kras mice treated with vehicle or IL33. **(O)** qRT-PCR for *Il33* expression in epithelial cells (Epit), tuft cells, CAFs and endothelial cells (Endot) isolated from Mist1-Kras-Dclk1-DTR-ZsGreen mice. **(P)** Immunofluorescence of pancreas from Mist1-Kras mice for IL33 and Vimentin (VIM). **(Q-R)** Immunofluorescence for AcTUB of pancreas from Mist1-Kras-Dclk1-DTR-ZsGreen mice treated with control IGG or anti-IL13 blocking antibody and related quantification. **(S)** qRT-PCR for the expression of *Sox9* in pancreas of Mist1-Kras-Dclk1-DTR-ZsGreen mice treated with control IGG or anti-IL13 blocking. Scale bars: 100 μm. Means ± SD. ∗: p ≤ 0.05; ∗∗: p ≤ 0.01; ∗∗∗: p ≤ 0.001.

**Figure S7.**
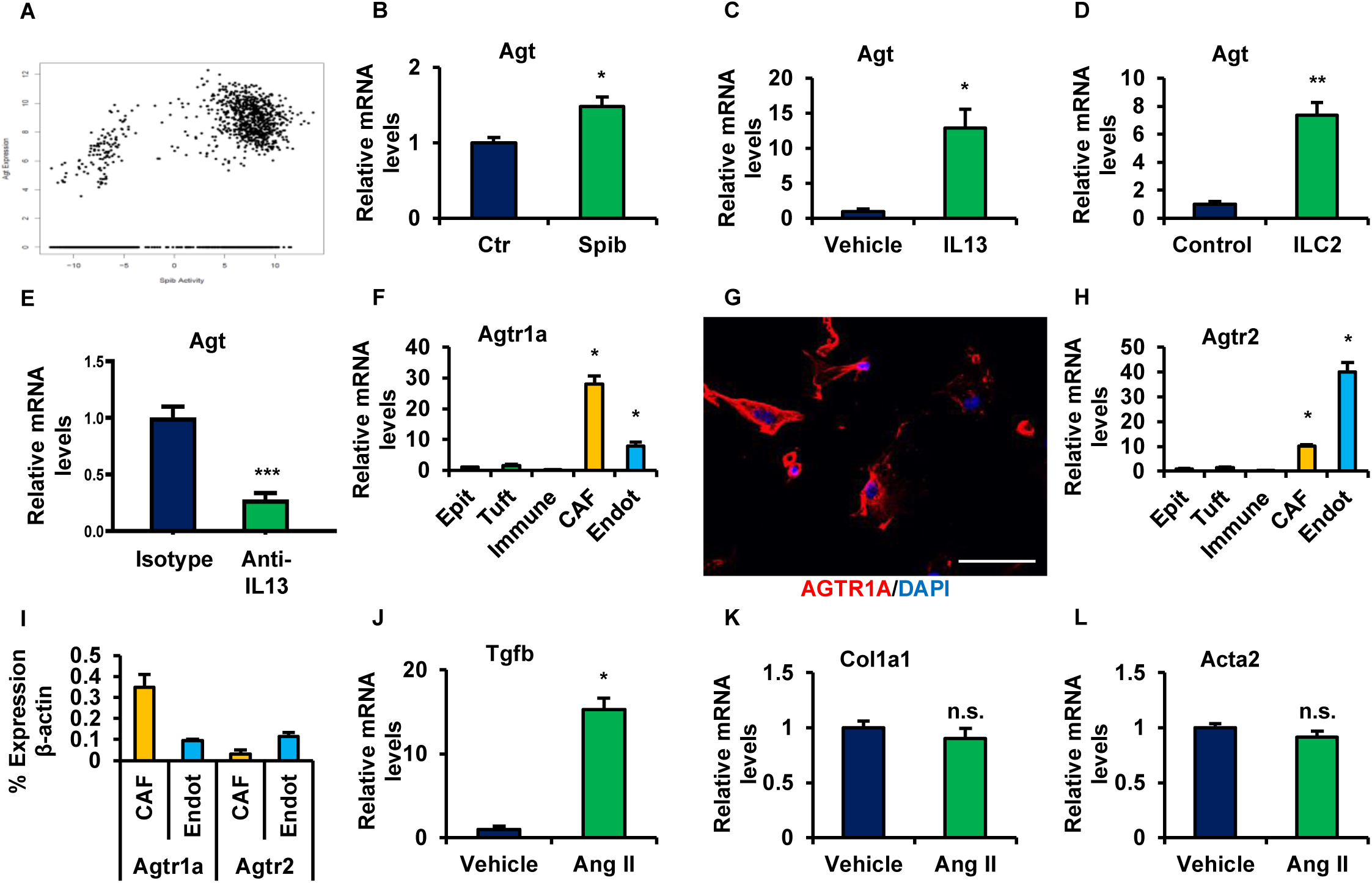
**(A)** Scatterplot showing the direct correlation between Agt expression and VIPER-inferred SPIB protein activity. **(B)** qRT-PCR for *Agt* expression in organoids generated from single cells isolated from pancreas of Kras mutant mice and transfected with control (Ctr) or *Spib*-expressing lentivirus (Spib). **(C)** qRT-PCR for *Agt* expression in organoids grown from single cells isolated from pancreas of Kras-Dclk1-DTR-ZsGreen mice in the presence of vehicle or IL13. **(D)** qRT-PCR for *Agt* expression in organoids grown from single cells isolated from pancreas of Kras-Dclk1-DTR-ZsGreen mice in the presence of control medium or ILC2. **(E)** qRT-PCR for the expression of *Agt* in pancreas of Mist1-Kras-Dclk1-DTR-ZsGreen mice treated with control IGG or anti-IL13 blocking antibody. **(F)** qRT-PCR for *Agtr1a* expression in epithelial cells (Epit), tuft cells, immune cells, CAFs and endothelial cells (Endot) isolated from Mist1-Kras-Dclk1-DTR-ZsGreen mice. **(G)** Immunofluorescence for AGTR1A of CAFs isolated from pancreas of Mist1-Kras mice. **(H)** qRT-PCR for *Agtr2* expression in epithelial cells (Epit), tuft cells, immune cells, CAFs and endothelial cells (Endot) isolated from Mist1-Kras-Dclk1-DTR-ZsGreen mice. **(I)** Relative expression of *Agtr1a* and *Agtr2* in CAFs and endothelial cells isolated from Mist1-Kras-Dclk1-DTR-ZsGreen mice. **(J-L)** qRT-PCR for the expression of *Tgfβ*, *Col1a1* and *Acta2* in CAFs isolated from Mist1-Kras-Dclk1-DTR-ZsGreen mice and treated with vehicle or angiotensin II (Ang II). Scale bars: 100 μm. Means ± SD. ∗: p ≤ 0.05; ∗∗: p ≤ 0.01; ∗∗∗: p ≤ 0.001.

## Notes

### Summary of Updates

We have updated the text by removing GEO accession numbers and data availability. Data will be made available after publication in a peer-reviewed journal.

